# S100A9 interacts with a dynamic region on CD14 to activate Toll-like receptor 4

**DOI:** 10.1101/2024.05.15.594416

**Authors:** Lauren O. Chisholm, Natalie M. Jaeger, Hannah E. Murawsky, Michael J. Harms

**Author notes:** These authors contributed equally to this article (Natalie M. Jaeger, Hannah E. Murawsky).

## Abstract

S100A9 is a Damage Associated Molecular Pattern (DAMP) that activates inflammatory pathways via Toll-like receptor 4 (TLR4). This activity plays important homeostatic roles in tissue repair, but can also contribute to inflammatory diseases. The mechanism of activation is unknown. Here, we follow up on a previous observation that the protein CD14 is an important co-receptor that enables S100A9 to activate TLR4. Using cell-based functional assays and a combination of mutations and pharmocological perturbations, we found that CD14 must be membrane bound to potentiate TLR4 activation by S100A9. Additionally, S100A9 is sensitive to inhibitors of pathways downstream of TLR4 internalization. Together, this suggests that S100A9 induces activity via CD14-dependent internalization of TLR4. We then used mutagenesis, structural modeling, and *in vitro* binding experiments to establish that S100A9 binds to CD14’s N-terminus in a region that overlaps with, but is not identical to, the region where CD14 binds its canonical ligand, lipopolysaccharide (LPS). In molecular dynamics simulations, this region of the protein is dynamic, allowing it to reorganize to recognize both S100A9 (a soluble protein) and LPS (a small hydrophobic molecule). Our work is the first attempt at a molecular characterization of the S100A9/CD14 interaction, bringing us one step closer to unraveling the full mechanism by which S100A9 activates TLR4/MD-2.

## INTRODUCTION

S100A9 is a small calcium-binding protein produced in copious quantities by neutrophils (1). When released into the extracellular space, it acts as a DAMP (Damage Associated Molecular Pattern) that activates pro-inflammatory innate immune pathways (2) (Figure 1A). S100A9 plays important roles in the response to tissue damage and related processes such as angiogenesis (3, 4). When dysregulated, however, it contributes to a variety of poor health outcomes including cancer (5–7), neurodegenerative disorders (8, 9), and chronic inflammatory diseases (10).

**Figure 1:**
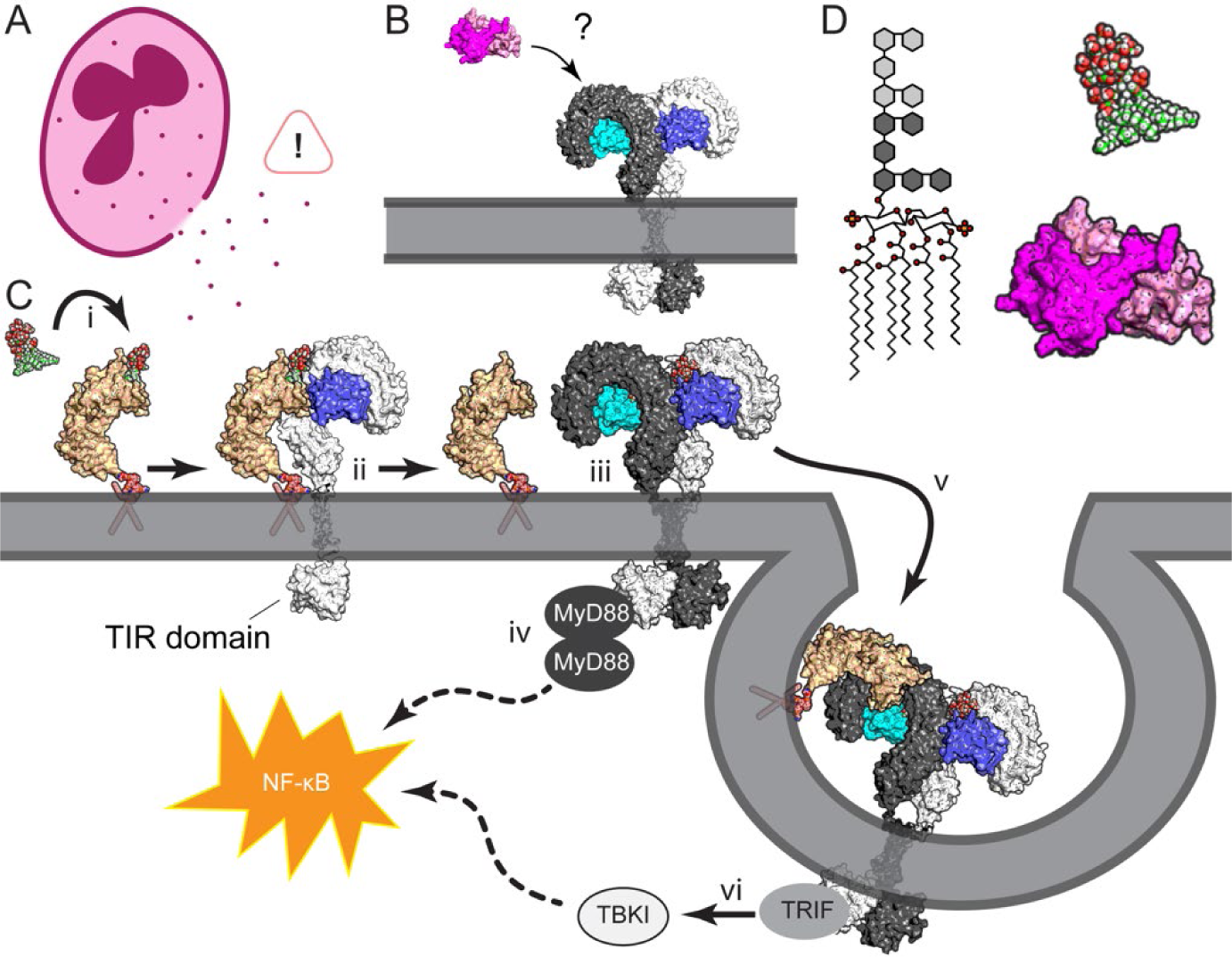
TLR4/MD2 responds to LPS and S100A9. A) S100A9 is released into the extracellular space by neutrophils, where it triggers inflammatory pathways. B) The mechanism by which the S100A9 dimer activates TLR4/MD-2 is unknown. The S100A9 structure is shown in pink/magenta: RCSB ID 1IRJ (33). The TLR4/MD-2 structure is gray/blue: 3FXI (34). C) In this scheme, lipopolysaccharide (LPS) is shown in spheres with green carbons; CD14 is shown in tan, with a modeled GPI linker in spheres. The CD14 structure is 4GLP (25). The PubChem CID for the GPI linker is 145996624 (35). TLR4 and MD-2 are shown as in B. In this mechanism, LPS binds to CD14 (i). CD14 then delivers LPS to a TLR4/MD-2 complex (ii), which homodimerizes with another TLR4/MD-2 heterodimer (iii). This brings the two TLR4 molecules’ TIR domains together, allowing them to bind MyD88, which activates a multi-step pathway (dashed lines) leading to activation of NF-κB (iv). Membrane-anchored CD14 and TLR4/MD-2 can then later internalize (v), activating the TRIF-dependent pathway which further amplifies the NF-κB response (vi). D) Schematic structure of *E. coli* LPS. Hexagons are sugar moieties in the inner and outer core; lines are carbons; red circles are oxygens. Space-filling models of LPS and S100A9 are shown on the same scale to the right.

S100A9 activates inflammation through Toll-like receptor 4 (TLR4) (3, 6, 11, 12); however, its mechanism of action remains poorly understood (Figure 1B). There is some evidence that S100A9 directly binds to TLR4 (11), and a handful of mutations to S100A9 and TLR4 are known to disrupt its activity in *in vitro* assays (13, 14). Developing a well-supported biochemical mechanism has, however, proved difficult. This lack of mechanistic understanding has had important consequences: a drug purportedly targeting S100A9 failed in phase III clinical trials due to lack of specificity (15).

Although it is unclear how S100A9 activates TLR4, the mechanism of action for TLR4’s canonical ligand is well understood (Figure 1C). TLR4 is known for responding to lipopolysaccharide (LPS), a glycosylated phospholipid from the outer membrane of gram-negative bacteria (16, 17). It does so in concert with the adapter protein MD-2, which accommodates the acyl chains of LPS in a large hydrophobic pocket (18). Binding of LPS promotes dimerization of TLR4, which then triggers the MyD88 inflammatory pathway via its intracellular TIR domain (16, 18–21). A third protein, CD14, plays an important supporting role by delivering LPS to the TLR4/MD-2 complex (22). CD14 binds LPS in an N-terminal pocket, shielding the acyl chains of LPS from solvent (23–27). It is anchored to the membrane by a C-terminal GPI anchor (22, 28, 29). In addition to delivering LPS, CD14 promotes internalization of the LPS-bound TLR4/MD-2 complex (30, 31), which triggers the TRIF-mediated inflammatory pathway (32).

TLR4/MD-2 and CD14 recognize LPS by its size and hydrophobicity, yet these features differ radically from S100A9 (Figure 1D). The portion of LPS recognized is ∼2 kDa—about ten times smaller than the 26 kDa S100A9 dimer. Furthermore, LPS has multiple hydrophobic acyl chains and is prone to micelle formation, while S100A9 is highly soluble in water. How does a soluble protein activate a receptor tuned to recognize a small hydrophobic moiety?

To gain a foothold to study this problem, we decided to focus on the interaction between CD14 and S100A9. In LPS signaling, CD14 is upstream of TLR4/MD-2, delivering LPS to the complex. We hypothesized that CD14 could be playing a similar role for S100A9. We reasoned that understanding the first step in the activation pathway would serve as a useful starting point for unraveling the total mechanism. In doing so, we are following up on previous observations that S100A9 and CD14 interact directly *in vitro*, as well as co-localize and co-internalize in immune cells (36). In line with this, we previously observed that TLR4, MD-2, and CD14 are all necessary for S100A9 to activate NF-κB signaling in a cell-based functional assay (37).

We set out to probe the role of CD14 in S100A9-mediated activation of TLR4/MD-2. We started with function: Does CD14 act as a delivery molecule for S100A9? If not, why does CD14 need to be present? We then turned to structure: What is the molecular basis for the S100A9/CD14 interaction? How does it compare to that of LPS? We tackled these questions using a combination of cell-based functional assays, site-directed mutagenesis, *in vitro* biochemical characterization, structural modeling, and molecular dynamics simulations. We found that the membrane anchor on CD14 was essential for activity with S100A9 and demonstrated that S100A9 signals more strongly through the TRIF pathway than LPS. This is consistent with CD14 promoting internalization of S100A9/TLR4 complexes, rather than behaving as a simple delivery molecule for S100A9. We also found that CD14 binds S100A9 and LPS at distinct, but overlapping, sites. The binding region of CD14 is highly dynamic, allowing it to take on different conformations and thus bind different molecules. This work is the first attempt at a molecular characterization of the S100A9/CD14 interaction, bringing us one step closer to unraveling the full mechanism by which S100A9 activates TLR4/MD-2.

## RESULTS

### CD14 improves the ability of S100A9 to activate the TLR4/MD-2 complex

We first set out to replicate the previous finding that CD14 was necessary for S100A9 to activate TLR4 (36, 37). To do so, we used an established *in vitro* functional assay in which we transiently transfect HEK293T cells with plasmids encoding human TLR4, MD-2, CD14, and an NF-κB luciferase reporter (13, 14, 37, 38). We then add potential agonists to the cell media and measure the expression of luciferase. HEK293T cells are excellent for such studies, as they natively express the required downstream effectors, but not TLR4, MD-2, or CD14 (39). The TLR4-induced NF-κB response induced by either S100A9 or LPS is thus strictly dependent on the presence of these transfected components (37). This allows us to study the relative contributions of each protein to the pro-inflammatory response with different agonists.

As a positive control for the assay, we first tested the ability of commercially available *E. coli* K-12 LPS to activate NF-κB signaling in the presence and absence of transfected CD14. As expected (22), we observed dose-dependent luciferase activity in the presence of CD14, but no activity its absence (Figure 2A).

**Figure 2:**
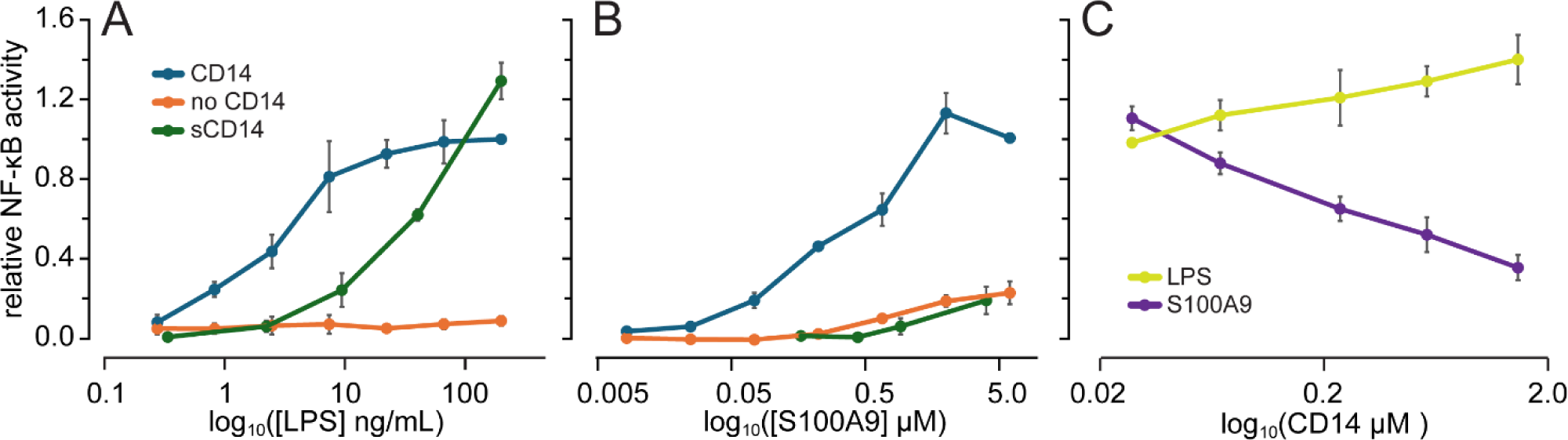
Membrane-anchored CD14 increases the response of TLR4/MD-2 to S100A9. A) NF-κB activity induced by increasing concentrations of purified LPS added to the growth media. The assay was done in HEK293T cells transiently transfected with human TLR4 and MD-2, +/− CD14. Series represent experiments done with CD14 transfected (blue), no CD14 transfected (orange), or purified soluble CD14 added to the growth media in the absence of transfected CD14 (green). Points are the means of at least three biological replicates. Error bars are standard error on the mean. B) NF-κB activity induced by increasing concentrations of purified human S100A9 to the growth media. Colors are the same as in panel A. C) Effect of adding increasing concentrations of purified soluble CD14 on the activity of cells transfected with CD14 in the presence of 20 ng/mL LPS (yellow) or 1 μM S100A9 (purple).

We next repeated this experiment using recombinantly purified human S100A9 as an agonist. As we have done previously, we included polymyxin B (PB) in our S100A9 experiments (13, 14, 37). PB sequesters LPS and prevents artifactual activation due to LPS contamination. As seen before (13, 14, 37), when we transfected plasmids encoding TLR4, MD-2, and CD14, we observed strong dose-dependent activation of NF-κB (Figure 2B). Transfecting only TLR4 and MD-2, but not CD14, dramatically lowered the response to S100A9. At 2 μM hS100A9, for example, NF-κB activity drops by 5-fold in the absence of CD14 (Figure 2B). (The presence of a small amount of S100A9 activity in the absence of CD14 is slightly different than one previous report (36), which found that CD14 was strictly required for S100A9 activity. This discrepancy could reflect a difference in the cell lines, as the previous experiments utilized THP1 and mouse BMDCs rather than HEK293T cells.)

### Soluble CD14 cannot replace membrane-bound CD14 for S100A9 signaling

We hypothesized that CD14 is delivering S100A9 to TLR4/MD-2, analogous to its function for LPS. To test this hypothesis, we took advantage of a soluble form of CD14 (sCD14). Although CD14 is typically produced with a GPI-linker that anchors it to the cell membrane, cells also export a form without this post-translational modification (40). sCD14 is present in the blood at ≥ 2 mg/L in healthy individuals, and is elevated during inflammation (41–43). Unanchored sCD14 scavenges extracellular LPS from the environment and transfers it to any cells expressing the TLR4/MD-2 complex (26, 40, 44, 45).

If CD14 is simply delivering S100A9 to TLR4/MD-2, we predicted that sCD14 would be able to replace anchored CD14 in our assay. To test this prediction, we expressed and purified sCD14 out of HEK293F cells. We then transfected HEK293T cells with plasmids encoding TLR4 and MD-2, but not CD14. In a separate tube, we pre-mixed purified sCD14 with either LPS or S100A9. We then applied these sCD14/agonist mixtures like other treatments and measured the luciferase response.

As a control, we first tested the ability of sCD14 to deliver LPS. Consistent with previous experiments (45, 46), sCD14 promoted LPS activation even in the absence of membrane-anchored (transfected) CD14 (green line, Figure 2A). We next tested the ability of sCD14 to deliver S100A9. Unlike LPS, sCD14 could not replace transfected CD14 for S100A9 activation (green line, Figure 2B). This result suggests that CD14’s role in S100A9 activity is more complex than a simple delivery/drop-off mechanism.

As both forms of CD14 are present biologically, we next tested the impact of sCD14 in cells with CD14 transfected, to determine whether the addition of non-productive sCD14 blocks the productive S100A9-CD14 interaction. As one might expect, adding sCD14 to LPS treatments improves TLR4 activity, as both forms of CD14 present can deliver LPS to TLR4 (yellow line, Figure 2C). Conversely, sCD14 inhibits S100A9 activation of TLR4 in a dose-dependent manner (purple line, Figure 2C). This implies that sCD14 either sequesters S100A9 and prevents it from binding to membrane-associated CD14, or that it binds to the TLR4/MD-2 complex non-productively in the presence of S100A9.

### Inhibition of TRIF dependent inflammation has a bigger effect on S100A9 than LPS-induced signaling

Our results indicate that CD14 plays a role in the S100A9 activation of TLR4 beyond simple delivery (Figure 2). To help explain this, we turned to what is known about LPS-induced inflammation. In this pathway, one of the roles of membrane-associated CD14—but not soluble CD14—is to promote internalization of TLR4/MD-2 (30–32). This activates the TRIF-dependent pro-inflammatory pathway in addition to the primary MyD88-dependent pathway (32) (Figure 1C). Although the TRIF pathway is typically associated with Type-I Interferon productions, both pathways result in NF-κB production (47), and thus are both measured by our HEK293T functional assay.

We hypothesized that internalization is important for S100A9 activity, explaining the difference between soluble and membrane associated CD14 activity on S100A9 activity. One prediction from this hypothesis is that inhibition of TRIF-dependent signaling would inhibit S100A9 activity. To test our hypothesis, we utilized our functional assay to measure NF-κB activity in response to S100A9 or LPS in the presence of the TRIF inhibitor MRT67307. This small molecule binds to and blocks the kinases TBKI and IKKε (48, 49), which operate downstream of TRIF (Figure 1C). We also tested the effect of the MyD88 inhibitor TJ-M2010-5 on activation by both ligands. This small molecule binds to the TIR domain of MyD88, blocking its homodimerization and thus ability to activate NF-κB (50, 51).

This experiment revealed a significant difference between S100A9 and LPS activation of TLR4 in the presence of MRT67307, but not TJ-M2010-5 (Figure 3). Because TRIF signaling requires internalization, this implies that activation by S100A9 triggers internalization of TLR4/MD-2. Taken together with our experiments using soluble CD14 (Figure 2), this suggests that at least part of CD14’s role in S100A9-induced activation of TLR4/MD-2 is promoting internalization of the complex. This aligns well with previous observations that S100A9 and CD14 co-internalize (36), and that a generic inhibitor of internalization, chloroquine, altered S100A9’s ability to activate of TLR4 (12).

**Figure 3.**
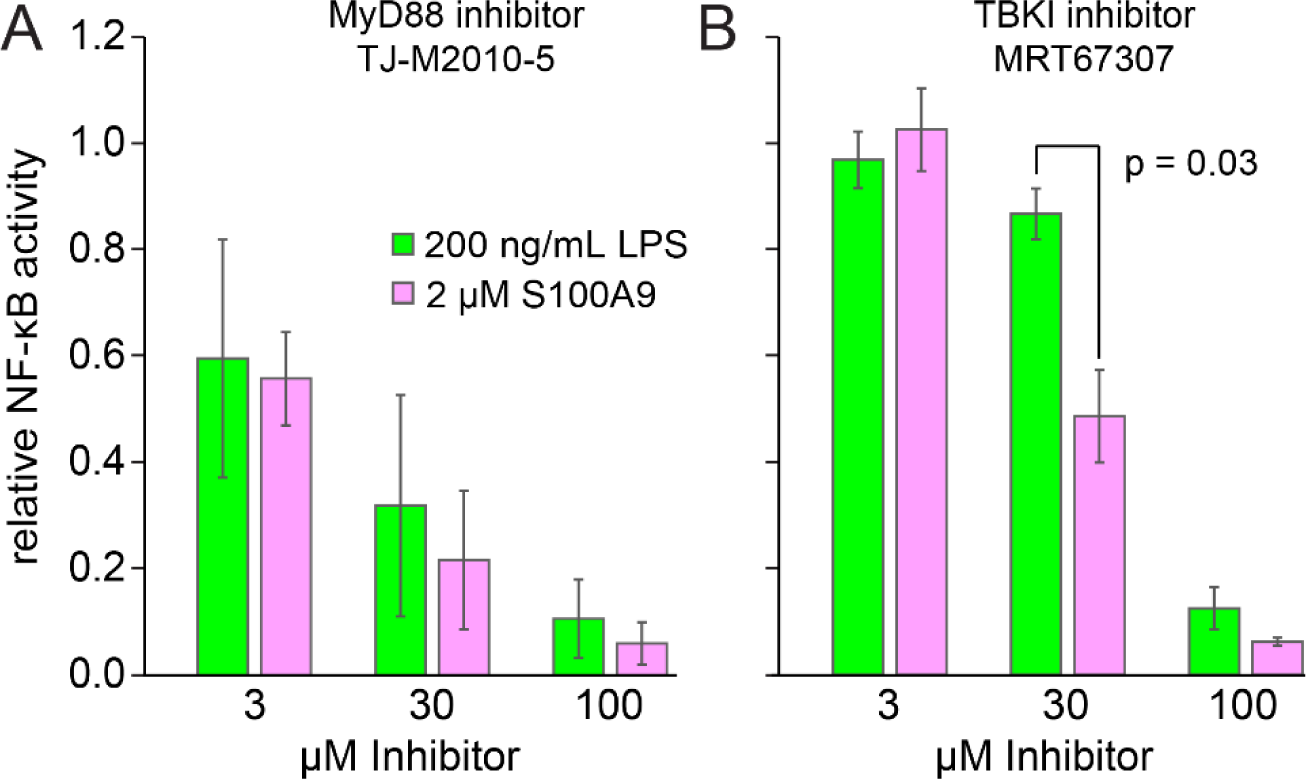
S100A9 activity may depend more on TRIF rather than MyD88 signaling. A) Effect of increasing concentrations of MyD88 inhibitor TJ-M2010-5 on NF-κB activity in response to 200 ng/mL LPS (green) or 2 μM S100A9 (pink). Bar heights are means of three biological replicates; error bars are standard errors on the mean. B) Effect of increasing concentrations of the TRIF inhibitor MRT67307 on activity induced by LPS or S100A9. Indicated p-value was calculated using a paired 2 sample students t-test. Colors are as in panel A.

### LPS and S100A9 interact with different residues of CD14

In addition to probing the function of CD14 in S100A9-dependent activation of TLR4/MD-2, we wanted to better understand the molecular basis for the interaction of S100A9 and CD14. Based on the different chemical structures of S100A9 and LPS (Figure 1) and the observed differences in S100A9 and LPS activation (Figures 2 and 3), we suspected that S100A9 and LPS interact differently with CD14, and that we could therefore separate their functions. To test this hypothesis, we studied the effects of an anti-CD14 antibody and mutagenesis to CD14 on LPS and S100A9 activity.

We first tested the ability of the anti-CD14 mAB MEM18 antibody to inhibit LPS and S100A9 activity in our assay. MEM18 is known to bind at amino acids 76-82 of human CD14 and block the CD14/LPS interaction (Figure 4A, yellow) (52). We co-treated cells with MEM18 and either LPS or S100A9. As expected, MEM18 inhibited LPS activation (p = 0.018), but had no effect on S100A9 activity (Figure 4B, 3C, yellow star). This result suggests that S100A9 interacts with a site on CD14 distinct from the LPS binding site.

**Figure 4:**
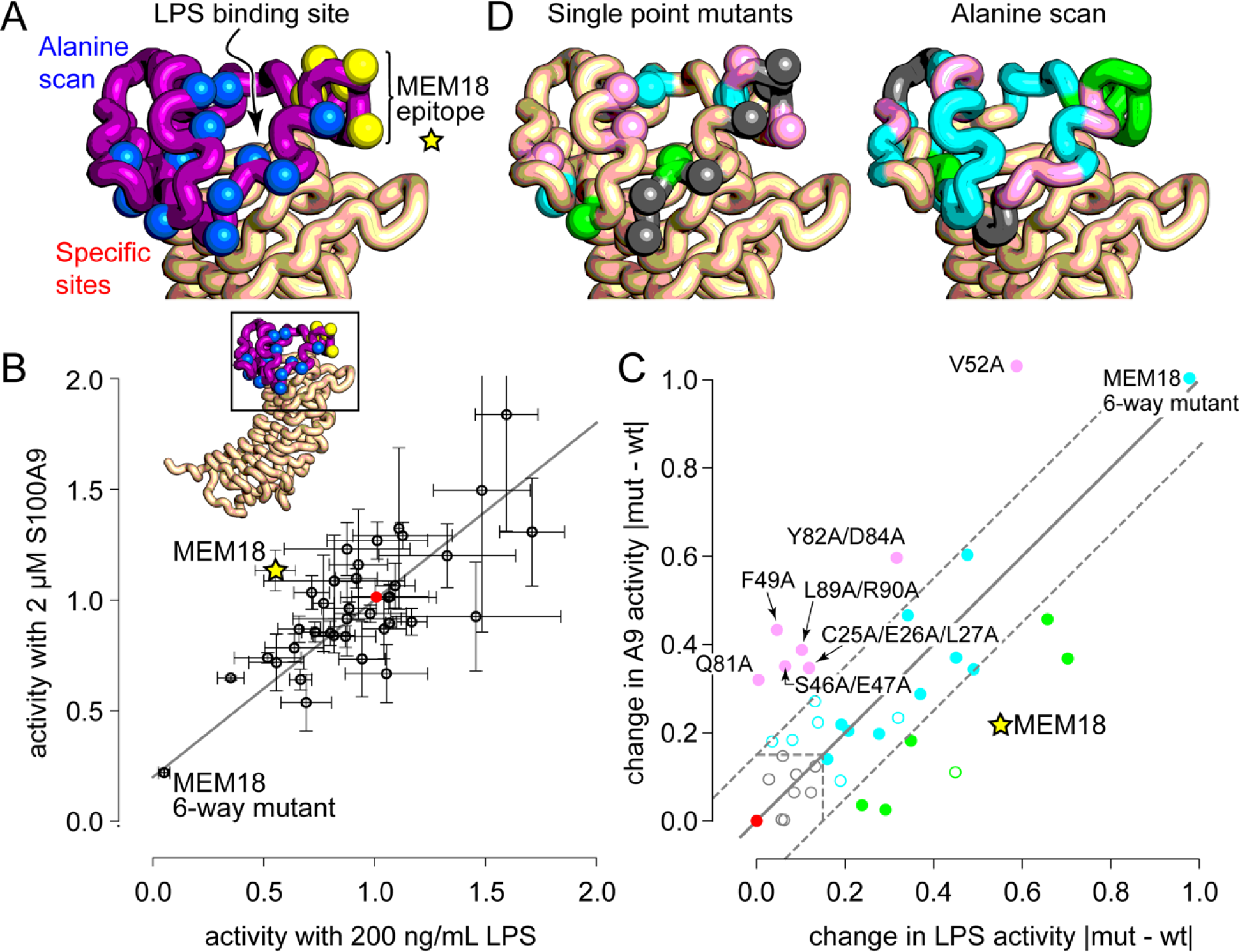
Mutations and antibody binding differentially affect LPS and S100A9 activity. A) Structure shows the region of CD14 under investigation. The purple region was probed with an alanine scan; the spheres are the C_β_ atoms of sites individually mutagenized; the yellow spheres are the known MEM18 epitope. B) Experimentally measured NF-κB activity in response to 200 ng/mL LPS or 2 μM S100A9 in cells with either wildtype CD14 and the MEM18 antibody (yellow star) or mutant versions of CD14 and no MEM18 (open circles). Points are the means of at least three biological replicates; error bars are standard errors on the mean. Activity is normalized to 1.0 for wildtype for both agonists (red circle). C) Magnitude of the change in LPS- and S100A9-induced NF-κB activity when antibody or CD14 mutations were introduced. This was calculated by the absolute value of the difference between the mutant and wildtype activity. Filled points have a mean difference at least two experimental standard deviations away from 0.0. Colors indicate effect type: larger effect on S100A9 (pink); larger effect on LPS (green); effect on both (blue); or no effect (gray). Wildtype CD14 is shown as a red point for reference. Mutations that had a larger effect on S100A9 are labeled. D-E) Results for single-mutants (D) and alanine scan (E) from panel C mapped onto the CD14 structure in the same orientation as in panel A. Colors match effect-type from panel C.

We next employed site-directed mutagenesis, using the four following strategies to select mutations. 1) We introduced point mutants that had been previously reported to alter LPS activity (23). 2) We selected amino acids near the LPS binding site based on the crystal structure of human CD14 (Figure 4A). This structure does not have LPS bound; however, the LPS binding pocket is known from previous biochemical experiments and docking studies (24, 25). 3) We mutagenized each of the sites in the MEM18 epitope individually and then all together. Finally, 4) We performed an alanine-scan of N-terminal residues 25-100 of CD14. This corresponds to the first 75 ordered amino acids in the protein (Figure 4A, purple). The first nineteen amino acids are a post-translationally removed signal peptide, while residues 20-25 are disordered in the structure. To maximize the effect size in the alanine scan, we introduced Ala mutations in blocks of three. For example, residues C25/E26/L27 were mutated to A25/A26/A27. We did not mutagenize existing alanine positions. For example, for the region A88/L89/R90, we introduced alanine at positions 89 and 90. For a list of all mutations we introduced, as well as their measured effects, see Table S1.

We used our functional assay to measure the NF-κB activity in response to 200 ng/mL LPS and 2 μM S100A9 for each of the CD14 mutants (Figure 4B). Many mutations had no effect, some moderately improved activity, and others moderately lowered activity. Only one set of mutations, converting every amino acid in the MEM18 epitope (residues 76-82) to alanine, completely disrupted activity. This did so for both LPS and S100A9, suggesting issues with CD14 expression or trafficking to the membrane.

We were interested in mutations that had differential effects on LPS and S100A9, so we calculated the magnitude (i.e. absolute value) of the effect of each mutant on LPS- or S100A9-induced activity. We focused on magnitude rather than sign to maximize our ability to identify sites where mutations modulated activity, rather than attempting to dissect deleterious or favorable effects. The results of this analysis are shown in Figure 4C-E. Most mutations had either no effect (gray points) or had similar effects on both LPS- and S100A9-induced activity (blue points). Seven CD14 mutations had a larger effect on S100A9 than LPS (pink points), while five mutations had a larger effect on LPS than S100A9 (green points).

We then plotted the effects of these mutations onto the crystal structure of human CD14 (Figure 4D-E). Broadly, mutations that specifically altered S100A9-induced activity occurred in two regions. The first region was perturbed by the single mutations F49A and V52A, as well as the alanine scan mutations to C25A/E26A/L27A and S46A/E47A. The second region was perturbed by D81A, as well as alanine mutations to Y82A/D84A and L89A/R90A.

These S100A9-specific mutations were scattered among mutations that had different effects: altering both LPS and S100A9, changing LPS only, or having no effect at all (Figure 4D). This suggests that LPS and S100A9 share at least partially overlapping binding sites. We also observed some peculiar combined effects within the mutations we tested. The L89A/R90A mutation disrupted S100A9 more than LPS, while the R90A single mutant disrupted LPS more than S100A9. Likewise, D81A disrupted S100A9 and had little effect on LPS, despite being part of the MEM18 epitope that disrupts LPS but not S100A9.

### A computational model predicts S100A9 interacts with CD14 via two interfaces

To try and make sense of these results, we used AlphaFold2 (53–55) to generate a model of an S100A9 dimer docked to a CD14 monomer (Figure 5A). The top-ranked docking model has S100A9 bound to the N-terminus of CD14 via two interfaces, marked in cyan (Surface I) and green (Surface II) on the structure (Figure 5A). These surfaces correspond approximately to the two regions that our mutant screen highlighted as important. Based on this model, we hypothesized that we observed only moderate effects for our tested mutants because these mutants left one or the other interface intact. We reasoned that we would need to disrupt both interfaces to completely disrupt the ability of CD14 to interact with S100A9.

**Figure 5.**
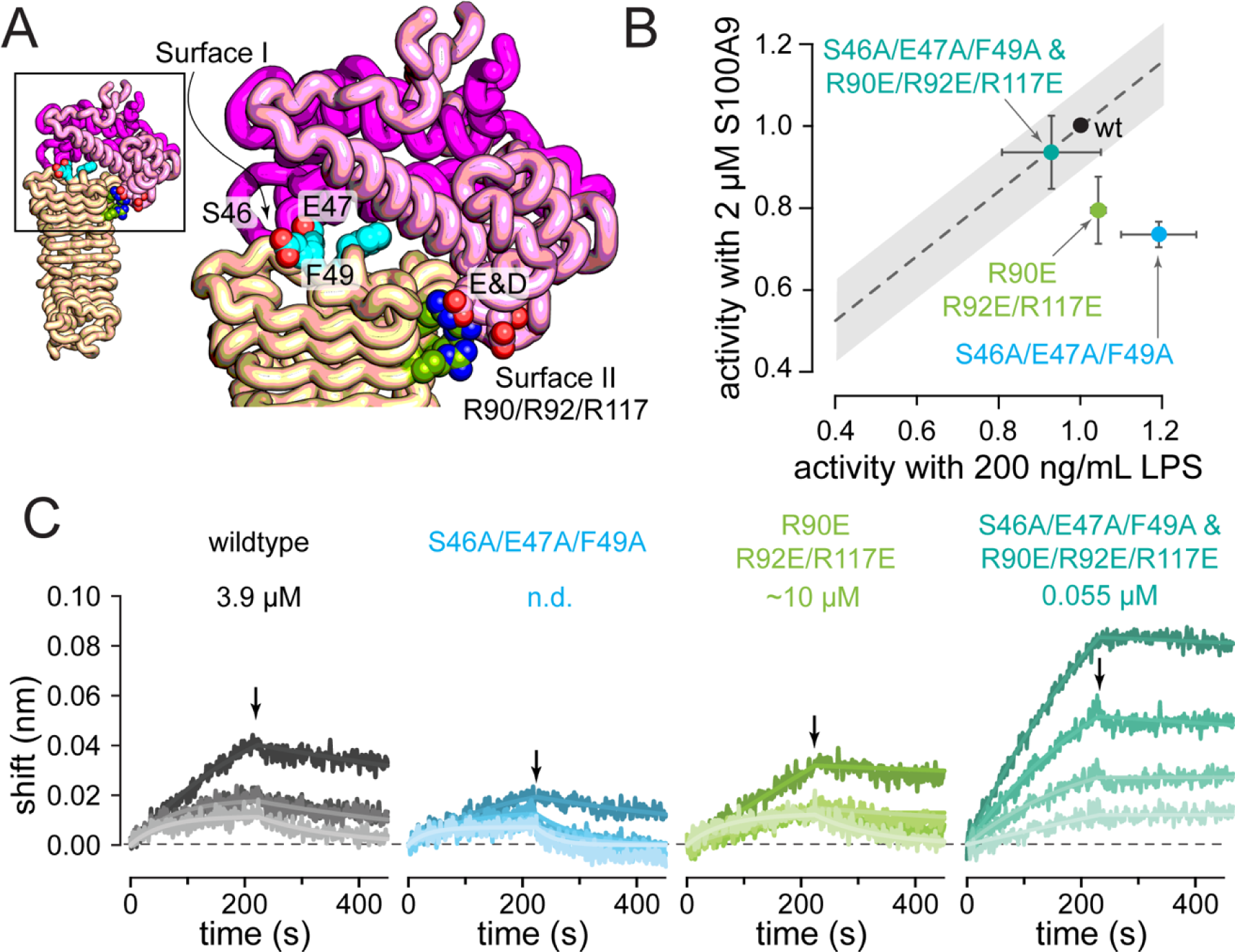
AlphaFold2 model predicts two binding interfaces on CD14. A) Structure showing the AlphaFold2 docking model. The S100A9 dimer is shown in pink (chain A) and magenta (chain B). CD14 is shown in tan. The residues we selected at the surface are shown as spheres: surface I is colored cyan, surface II is colored green. B) Experimentally measured NF-κB activity in response to 200 ng/mL LPS or 2 μM S100A9 in cells with wildtype CD14 and the indicated mutants of surface I, surface II, or surface I & II mutants. Points are the means of at least three biological replicates; error bars are standard errors on the mean. Activity is normalized to 1.0 for wildtype for both agonists (black circle). C) Bio-Layer Interferometry binding data for S100A9 interacting with sCD14. 50 nM biotinylated S100A9 was immobilized on a streptavidin sensor. Traces show response to four different concentrations of sCD14 (0.5μM, 1 μM, 2 μM, and 4 μM). Small arrow on each panel shows the point on that experiment where we switched from sCD14 to buffer. Model fits are shown as lines. Plots are representative from one biological replicate. Our best estimate of the S100A9/CD14 K_D_ for each variant are shown above each plot (see text).

To test the two-interface docking model, we generated mutants of CD14 expected to disrupt either one or both surfaces. For Surface I, we noted that the wildtype sequence of CD14 contains bulky hydrophobic residues (W45, F49, etc.). Therefore, we chose to take an existing alanine mutant with a deleterious effect on S100A9 activation—S46A/E47A—and expand it to include the individually deleterious mutant F49A. For Surface II, we noted that the CD14 has many positively charged residues in this region. Specifically, three arginine residues (R90, R92, and R117) were in the vicinity of negatively charged amino acids on S100A9. We also had some evidence for the importance of this region, as the alanine mutant L89A/R90A improved S100A9 activity. To attempt to disrupt this interface, we introduced a negatively charged glutamate in place of arginine at all three of these positions.

Based on this reasoning, we generated three CD14 mutants: Surface I (S46A/E47A/F49A), Surface II (R90E/R92E/R117E), and Surface I & II (S46A/E47A/F49A & R90E/R92E/R117E). We tested the ability of these CD14 mutants to support both LPS and S100A9 activation of TLR4 (Figure 5B). As predicted, we found that the Surface I mutant lowered S100A9 activity by 30% (p = 0.007). It also slightly increased LPS activity (p = 0.078). Surface II mutant decreased S100A9 sensitivity by 20% (p = 0.06), and had no measurable effect on the LPS response. We next tested our combined Surface I & II mutant. To our surprise, the mutant was indistinguishable from wildtype CD14 when stimulated by either S100A9 or LPS (Figure 5B). The combined effect of two deleterious mutations was to have no effect at all.

One explanation of this observation is that these mutations are disrupting some feature of CD14 besides its ability to interact with S100A9. We know these proteins are being expressed and trafficked properly, as they promote activity with LPS. We wanted, however, to directly probe the hypothesis that CD14 and S100A9 interact at these sites, and that these mutations modulate binding.

To determine if these mutations altered binding, we measured the binding of wildtype CD14 and the three mutants to wildtype S100A9 using bio-layer interferometry (BLI). We biotinylated S100A9 and immobilized it on streptavidin sensors. We validated our experimental conditions with an anti-S100A9 mAb, measuring its K_D_ as 0.15 nM (Figure S1). We then measured soluble CD14 binding at concentrations between 0.5 µM and 4 µM. Despite extensive work to optimize the blocking and binding conditions, we struggled to collect high quality data for these proteins. Our traces were often barely above background. (The antibody, by contrast, gave excellent results; Figure S1). This is consistent with previously published surface plasmon resonance studies (36), which found that the CD14-S100A9 interaction gave low signal. Given these difficulties, we applied a quality-control filter to all experiments, excluding any experiment for which the regressed binding model had R^2^ < 0.95 when compared to the experimental data.

We started by measuring the interaction between S100A9 and wildtype sCD14. Of the four bio-replicates we attempted, only two yielded data that passed quality control. The mean K_D_ for these two replicates was 3.9 μM, with a 95% confidence interval of 0.2 to 80 μM (Figure 5C). This value is consistent with the previously measured K_D_ of 0.2 μM, accounting for different conditions and experimental methodologies (36).

We next introduced the Surface I and Surface II mutants and re-measured binding. The signal was even weaker for these proteins than for the wildtype protein: we could only reliably fit a model to one of our six biological replicates. The Surface I mutant yielded no fittable experiments (Figure 5D), while the Surface II mutant yielded one fittable experiment, with a K_D_ of 10 μM (Figure 5E). This apparent disruption of binding—manifesting as low BLI signal—is consistent with our functional assays, which found that both individual surface mutants lowered activity with S100A9 (Figure 5B). Intriguingly, when we combined the two surface mutants, we obtained much higher binding signal, with all three biological replicates yielding measurable signal (Figure 5E). The K_D_ for these three replicates was 55 nM, with a 95% confidence interval of 2 to 1000 nM.

Taken together, these mutant studies support the basic features of the docking model. Mutations at surface I and II perturb activity and binding. Importantly, the effects of the mutations track across both activity and binding: the mutants with decreased binding have lower activity, while the double mutant with increased binding recovers activity. At the same time, our simple reasoning based on charge and size complementarity was not able to predict the effect of mutations at both surfaces.

### MD simulations reveal the N-terminus of CD14 is dynamic

To better understand the molecular origins of these complicated patterns of mutational effects, we turned to atomistic molecular dynamics (MD) simulations. These simulations allowed us to relax the docking model, as well as compare the contacts made between CD14 and S100A9 with those seen for CD14 and LPS. We simulated CD14 alone, CD14 interacting with LPS, and CD14 interacting with S100A9. (See the methods for a detailed description of how we constructed our models and ran the calculations.) We ran three or four 500 ns simulations per condition, giving 1.5-2.0 μs of total simulation time for each.

We observed that the region of CD14 that binds to LPS and S100A9 was dynamic in all simulations. One of the consistent differences was the position of the “lid” formed by residues 25-55 (Figure 6A-C, yellow). In simulations of CD14 alone, this region collapsed into CD14’s binding pocket, burying lid residues W45 and F49 (Figure 6A). This closed conformation shields the hydrophobic residues of the LPS binding pocket from water molecules. This conformation differs from the crystal structure of apo CD14, which has the hydrophobic LPS binding pocket open and exposed to water (25). The conformation in the crystal structure is likely an artifact due to crystal-contacts stabilizing the open conformation (Figure S2).

**Figure 6.**
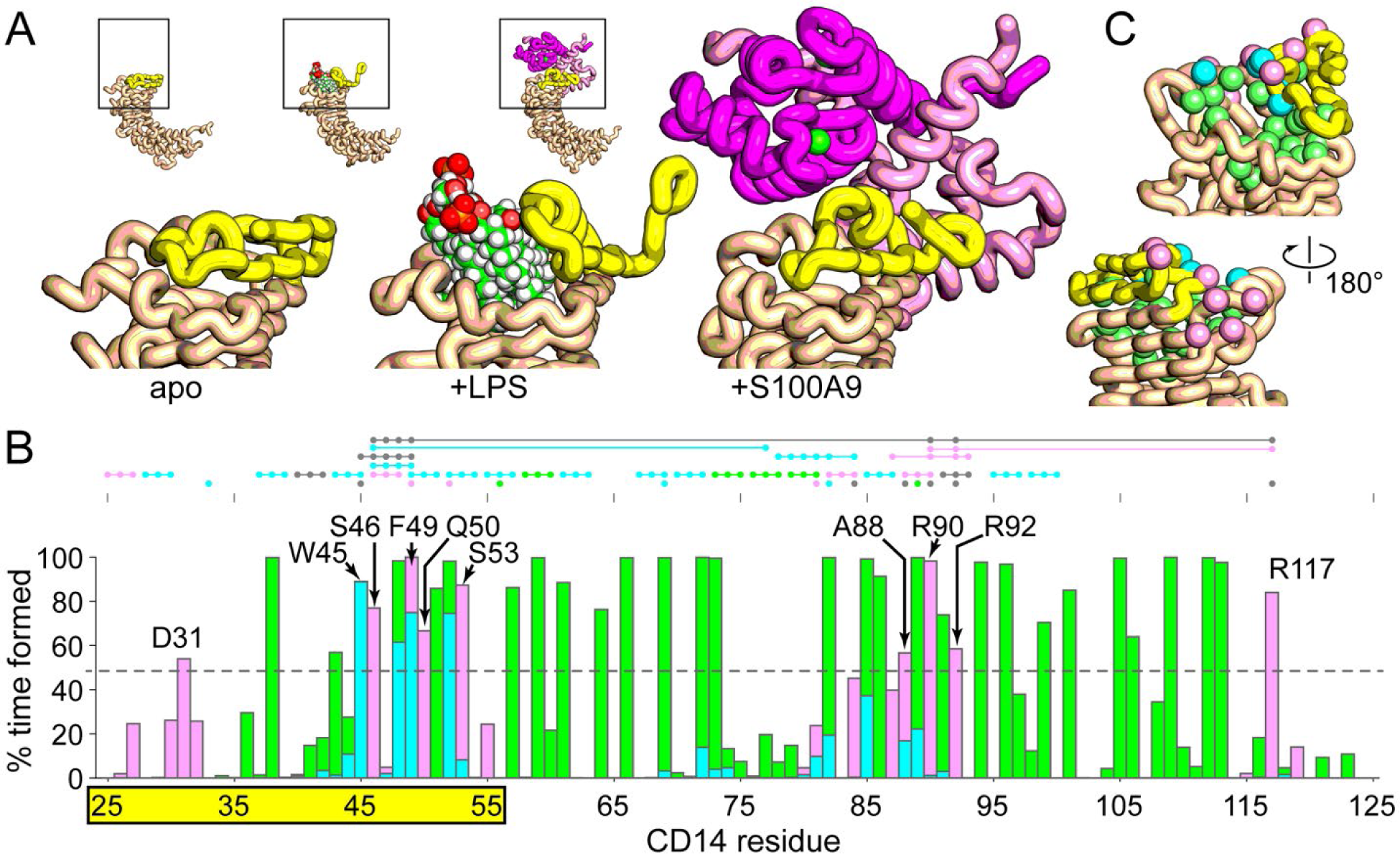
LPS and S100A9 interact with overlapping residues on CD14. A) Representative snapshots taken from simulations of CD14 with no ligand (left), LPS (middle), or S100A9 (right). The mobile lid (residues 25-55) is shown in yellow. S100A9 residues 1-3 and 91-114 are disordered and omitted for clarity. B) Bar plot shows the fraction of time each CD14 residue has at least one heavy atom within 4 Å of a heavy atom in S100A9 (pink) or LPS (green) across all simulations. Overlapping bars appear as blue. Residues participating in S100A9 interactions more than 50% of the time, but not LPS interactions, are annotated. Mobile lid residues W45 and F49 are also indicated, with the full span of the lid highlighted in yellow below the plot. Dots above the plot summarize the results from Figure 4C. Each point is a mutated residue; connected points indicate clones with multiple mutations. The color indicates the effect of that mutation from Figure 4C, where the mutation affects activation by: S100A9 alone (pink), LPS alone (green), both agonists (blue), or neither agonist (gray). C) Contacts populated more than 50% of the time in Figure 6B mapped onto the crystal structure of CD14. Color indicates the sorts of contacts made: S100A9 only (pink), LPS only (green), or both (blue).

In simulations with LPS, the aliphatic carbons of the LPS acyl chains interact with the hydrophobic binding pocket (Figure 6A). To accommodate the LPS molecule, the lid residues are displaced from the pocket. The lid residues noted above, W45 and F49, switch from interacting with hydrophobic residues in the binding pocket to interacting with the LPS acyl chains. This secures the LPS within the pocket and excludes water from interactions with hydrophobic regions of either LPS or the binding pocket. The CD14/LPS interaction is almost entirely hydrophobic in nature. We compared simulations of LPS and CD14 alone to the simulations of the CD14/LPS complex, finding that LPS binding buries an average of 1360 Å^2^ of nonpolar surface, but only 200 Å^2^ of polar surface.

In simulations with S100A9, CD14 interacts with similar, but distinct, residues when compared to LPS (Figure 6A). The CD14 lid residue F49 is in continuous contact with S100A9 over the course of the simulations, interacting with a hydrophobic patch formed by S100A9 residues M81, A84, W88, and the greasy aliphatic carbons of R85. In contrast, lid residue W45 bridges the LPS binding pocket and S100A9, interacting with both throughout the simulation. As one might expect for a protein-protein interaction, the CD14/S100A9 interface has more diverse interactions than the CD14/LPS interface. The interaction buries 880 Å^2^ and 730 Å^2^ of nonpolar and polar surface, respectively, reflecting both hydrophobic and hydrogen-bonding interactions across the interface.

A quantitative comparison of the CD14 residues in contact with LPS and S100A9 reveals the two agonists interact with distinct, but overlapping, sets of residues on CD14. Figure 6B shows the percent of time at least one atom from each residue of CD14 is in contact with LPS or S100A9. For simplicity, we defined any interaction formed more than 50% of the time as a contact. We found that LPS was in contact with 29 residues on CD14 (Figure 6B). Of these, 25 were specific to LPS, and not S100A9. As one might expect, these LPS-specific contacts are deep in the hydrophobic binding pocket of LPS (Figure 6C). S100A9, in contrast, only contacts 12 residues. Eight of these are specific to S100A9. These S100A9-specific contacts cluster outside the LPS binding pocket, both within the lid (D31, S46, Q50, and S53), as well as adjacent to the lid in the structure (A88, R90, R92, R117) (Figure 6C). These correspond closely to Surface I and Surface II from our binding experiments.

Finally, there are four residues, all in the lid region, that contact both S100A9 and LPS: W45, A48, F49, and V52. These residues take on different conformations and interactions depending on the ligand bound. When LPS is bound, they interact with acyl chains; when S100A9 is bound, they interact with a hydrophobic patch on the protein; when nothing is bound, they interact with the hydrophobic pocket of CD14.

### S100A9 binds to CD14 with competing binding modes

Visual inspection of the simulations also revealed that the relative orientations and contacts between S100A9 and CD14 fluctuated over the course of the simulations (Figure S3). To explore this phenomenon further, we clustered simulation frames based on features describing the orientation and contacts of the two proteins (see methods for details). Briefly, we recorded which residues in CD14 and S100A9 had heavy atoms in contact, which residues participated in hydrogen bonds, and the rotation matrix necessary to align S100A9 from a given frame to the starting conformation. This yielded 228 features per frame. We then sampled frames every 0.5 ns and used k-means to identify clusters of conformations with shared features. This revealed two conformational clusters.

Figure 7A and B show representative snapshots taken from the two clusters we identified. In the one cluster—the “open” conformation—residues 25-55 and 71-89 of CD14 move apart, exposing the hydrophobic pocket used to bind LPS. S100A9 forms numerous contacts with CD14 in this conformation (black and blue spheres, Figure 7A). In the other cluster—the “closed” conformation—CD14 residues 25-55 and 71-89 interact with one another, sequestering the hydrophobic pocket from solvent. This conformation loses many of the S100A9 contacts seen in the open conformation (blue spheres) but gains a handful of new S100A9 contacts on the opposite side of the protein (orange arrow, Figure 7B).

**Figure 7.**
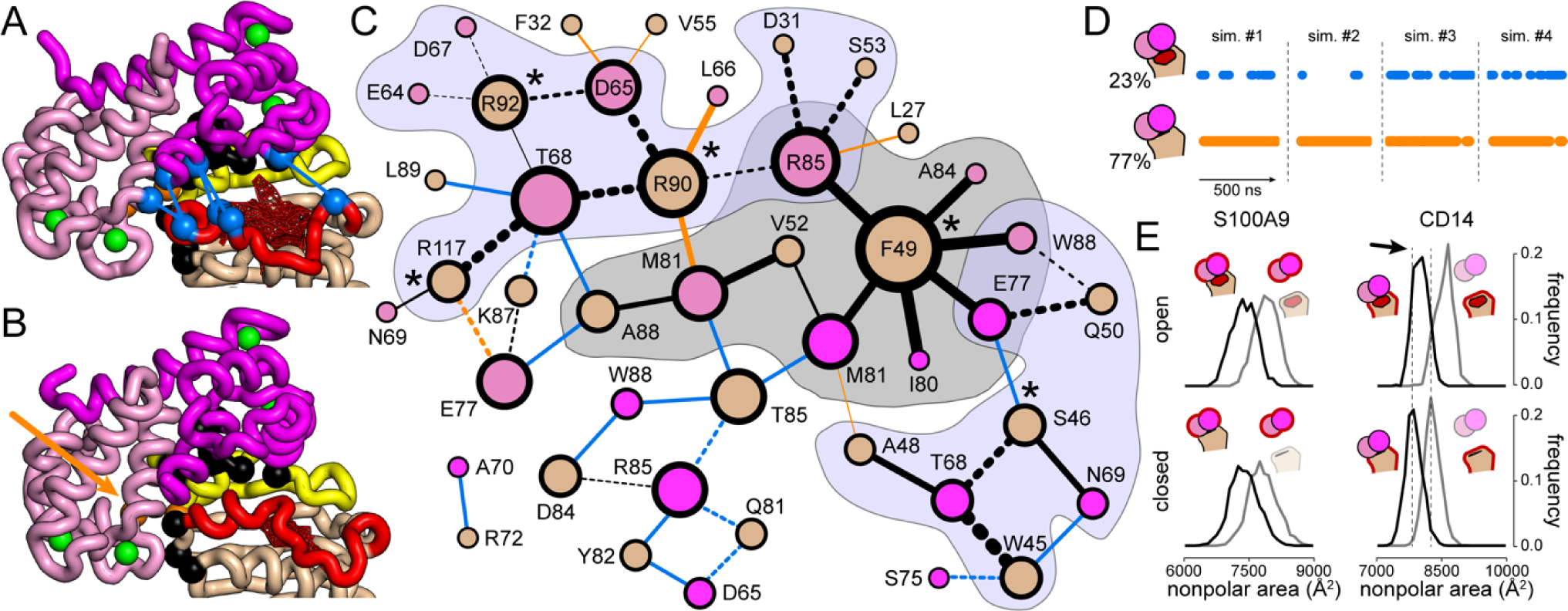
The S100A9/CD14 interface consists of competing interactions. A & B) Representative frames of the open (panel A) and closed (panel B) conformations of CD14 bound to S100A9. The S100A9 dimer is shown in pink (chain A) and magenta (chain B), with calcium ions shown as green spheres. Disordered S100A9 residues 1-3 and 91-114 are not shown for clarity. CD14 is shown in tan, with mobile regions shown in yellow (residues 25-55) and red (residues 71-89). Exposed hydrophobic surface in the binding pocket is shown as red mesh. The C_α_ atoms of residues participating in contacts between CD14 and S100A9 are shown in spheres whose colors indicate whether the contact is formed in both conformations (black), only open (blue), or only closed (orange). The closed-only contacts are on the opposite side of the structure, indicated with an orange arrow. C) Network of contacts formed between S100A9 and CD14 over the simulations. Nodes are amino acids in contact, with the color indicating the protein as in panels A & B. Node size indicates the number of contacts for that residue. Edges represent contacts. Edge width indicates the fraction of simulation time the interaction is formed from 10% (thinnest) to >90% (thickest). Edge color indicates whether the interaction is found in both conformations (black), specific to open (blue), or specific to closed (orange). A solid line is a non-polar contact; a dashed line is a polar contact. The gray region indicates the central hydrophobic surface between the molecules; the slate regions indicate adjacent networks of polar interactions. “*” icons indicate residues mutated in Figure 5. D) Plots show fluctuation between conformations over four replicate, 500 ns simulations. Points are frames where the proteins are in either the open (blue) or closed (orange) cluster. E) Nonpolar surface area for S100A9 (left) or CD14 (right) in simulation snapshots (black) or snapshots with the partner deleted (gray). The top row shows the results for the open conformation, the bottom row for the closed conformation. The shift in the black and gray distributions for CD14 (dashed lines) shows that the open conformation of CD14 has more exposed nonpolar surface than the open conformation.

The nature of the rearranged contacts can be seen by drawing the contacts as a network (Figure 7C). Notably, a central set of contacts is shared by both the open and closed conformations. CD14 residues F49, V52, and A88, form a central non-polar surface with carbons from S100A9 residues E77, I80, M81, A84, R85, and W88 (Figure 7C). This is surrounded by two networks of peripheral, largely polar, interactions. One network is built around CD14 residues W46, S46, and Q50, the other around CD14 residues R90, R92, and R117. These correspond to Surface I and II from Figure 5.

The open and closed conformations differ significantly in the number and nature of contacts on top of this shared core. The open conformation creates a whole new interface with residues 72-85 (blue spheres, Figure 7A; blue edges Figure 7C). This is adjacent to and complements the core interface formed between CD14 and S100A9. The closed conformation, by contrast forms only a few new contacts relative to the core network. These interactions tend to tie into existing core residues (for example, CD14 L27 interacting with S100A9 R85), or add new contacts between residues already in the network (for example, CD14 R90 with S100A9 M81).

Based on a naïve count of contacts, we would expect the open conformation to be favored over the closed conformation. It possesses 17 extra contacts, beyond the core network, between residues on S100A9 and CD14. When we studied the relative population of the two conformations over the course of the simulations; however, we found the opposite. Both conformations are populated in all four 500 ns simulations; however, the closed conformation is found about 3 times more often than the open conformation (77% versus 23%, Figure 7D). The bias towards the closed conformation is likely due to the exposure of hydrophobic surface area in the CD14 binding pocket in the open conformation (Figure 7A). This can be seen quantitatively in the amount of surface area buried when S100A9 and CD14 interact in either the open or closed conformations (Figure 7E). S100A9 buries similar amounts of nonpolar surface when it interacts with CD14 in either the open or closed conformations. By contrast, the open form of CD14, by itself, more exposed hydrophobic surface than the closed form (Figure 7E).

The S100A9/CD14 interface is thus complex. Contacts between CD14 and S100A9 favor the open form of CD14, while the tendency to bury hydrophobic surface favors the closed form of CD14. As a result, the S100A9/CD14 complex fluctuates between the two conformations over the course of the simulation.

## DISCUSSION

We set to do an initial molecular characterization of the interaction between CD14 and S100A9. Using *in vitro* functional assays, we found that CD14 dramatically improves the activation of TLR4/MD-2 by S100A9. We also found significant differences between membrane-anchored and soluble CD14. Extensive mutagenesis and computational studies revealed that S100A9 interacts with the N-terminus of CD14, at a site distinct from the LPS interaction site. This region appears to be dynamic, allowing identical residues to participate in interactions with both lipid acyl chains and a soluble protein. This work is an important step towards understanding the mechanism by which S100A9 activates TLR4.

### Proposed activation model

Our findings allow us to construct a model for how CD14 increases S100A9’s ability to activate TLR4/MD-2. The key observations informing this model are: 1) Membrane-anchored CD14 is necessary for potent S100A9 activity (Figure 2B). 2) Soluble CD14 inhibits activation of TLR4 by S100A9 in the presence of membrane-anchored CD14 (Figure 2C). 3) S100A9 signals through TRIF more than LPS does in these assays (Figure 3). 4) S100A9 interacts directly with CD14 (Figures 4 and 5).

These observations rule out a simple delivery mechanism in which CD14 drops off S100A9 to the TLR4/MD-2 complex (Figure 8A). One model that accounts for these observations has S100A9 binding directly to CD14, which then promotes internalization of TLR4/MD-2 (Figure 8B). This model is also consistent with existing work showing that blocking internalization differentially inhibits S100A9 activation of TLR4 (12), and that S100A9 and CD14 co-internalize (36). In the context of this model, the simplest explanation for the inhibitory activity of sCD14 would be that sCD14 sequesters S100A9 and prevents the productive CD14-S100A9 interaction (Figure 8C).

**Figure 8.**
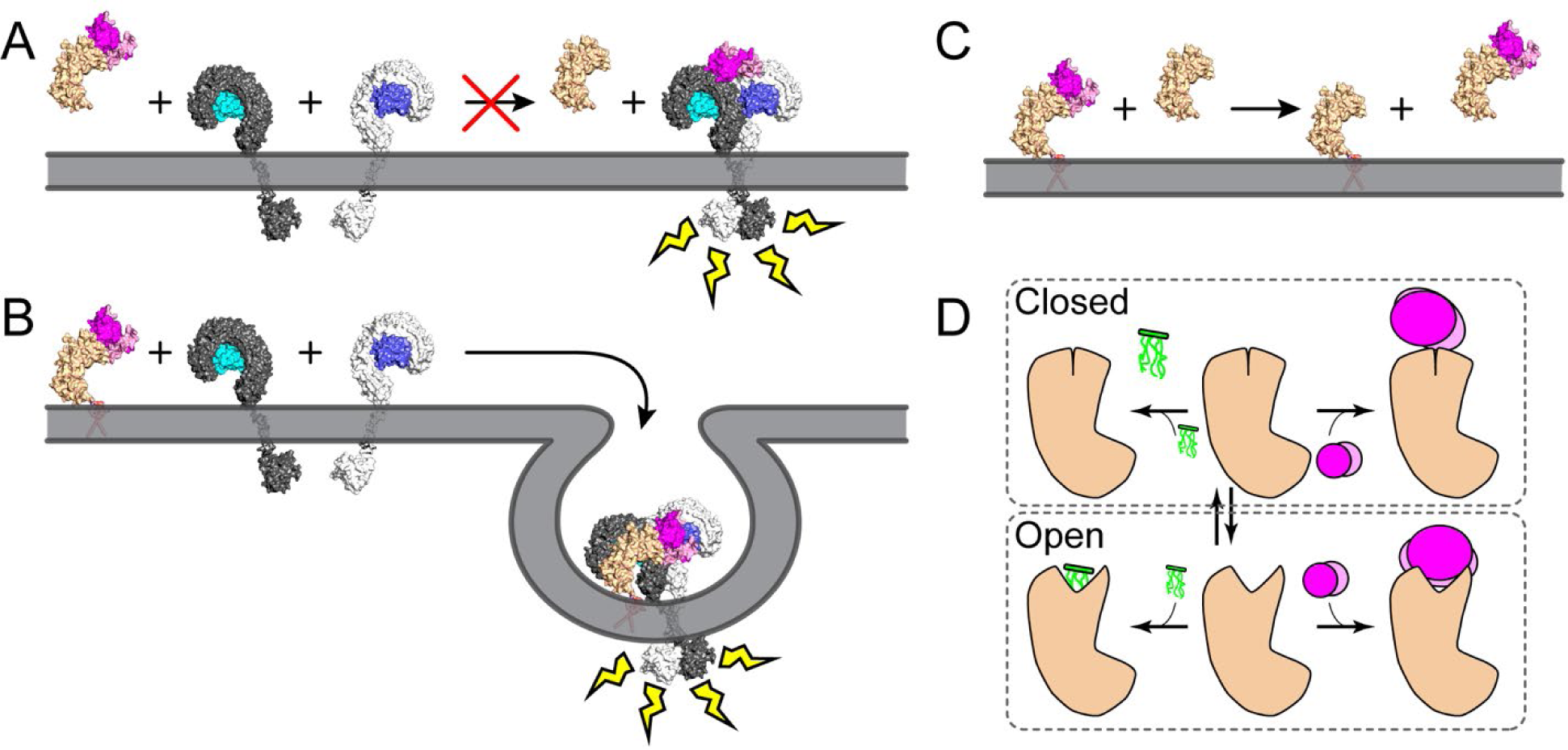
Models for the S100A9/CD14 interaction consistent with our data. A) Soluble CD14 (tan) cannot deliver S100A9 (pink) to TLR4/MD-2 (gray/blue) and trigger dimerization and activity. B) Membrane associated CD14 assembles a complex with TLR4/MD-2 and S100A9 that internalizes and triggers pro-inflammatory pathways. C) Soluble CD14 binds to and sequesters S100A9 away from membrane-associated CD14, thus preventing its activity. D) Schematic showing how CD14 binds to its ligands. CD14 (tan) is in equilibrium between a closed and open conformation. LPS (green molecule) binds exclusively to the open conformation; S100A9 (pink circles) can bind to either form in slightly different conformations.

We believe the model shown in Figure 8A-C is the simplest model that accounts for our functional observations. That said, even if this model is correct in broad outline, many questions remain. One of the most important questions is whether CD14/S100A9/TLR4/MD-2 assemble into a stable complex that is then internalized, or whether the interaction between TLR4/MD-2 and S100A9/CD14 is transient. Our data hint at the answer to this question. We found that weakening S100A9 binding is disruptive to CD14 function (Figure 5, mutating surfaces I and II alone), but strengthening the S100A9/CD14 interaction has no effect on activity (Figure 5, mutating surfaces I and II together). This result is consistent with quaternary complex formation; CD14 doesn’t need to “let go” of S100A9 after the initial binding event. Formation of this complex may even promote internalization. This is, however, speculative: further work is required.

Aspects of this model may also be wrong while the overall account remains correct. Our proposed mechanism by which sCD14 inhibits S100A9’s ability to activate is plausible, but speculative. While sCD14 could compete for S100A9 as shown in Figure 8C, it could also compete with binding sites on TLR4/MD-2, inhibiting the interaction with membrane associated CD14.

Finally, it is possible that the model is mostly wrong. For example, it is possible that CD14 does not need to directly interact with TLR4/MD-2 to promote activation, but instead must be present to allow S100A9 to promote internalization. This could be achieved if S100A9 does not bind to TLR4/MD-2 at all, but instead interacts with and reorganizes the membrane. In this case, the role of CD14 could be to nucleate lipid rafts in the vicinity of TLR4/MD-2 via its GPI-anchor. We favor a simple binding model because changes in S100A9/CD14 binding affinity correlate with changes in S100A9 activity, suggesting a role for binding (Figure 5). S100A9 is also known to interact directly with TLR4/MD-2, with a reported K_D_ of 3 nM (11), also favoring a binding model. Despite these hints, a non-binding mechanism has not been fully ruled out by our data or other reports in the literature.

### Biological implications

Regardless of the precise mechanism of action, our results have three important biological implications. The first is that S100A9 can likely activate a slightly different set of downstream pathways than LPS. The MyD88 and TRIF pathways are temporally distinct and produce different inflammatory markers (47). Indeed, other TLR4 agonists have been shown to induce differential activation of TRIF vs MyD88 dependent pathways (56, 57) as a method of modulating the immune response. Because S100A9 appears to signal more strongly via TRIF than MyD88, it may activate different outputs than LPS.

The second implication is that only a subset of TLR4+ cells are likely sensitive to S100A9. This is because not all cells that express TLR4/MD-2 also express membrane-anchored CD14 (22). Indeed, one of the roles of sCD14 is to bring LPS to cells that do not express the membrane anchored form. We predict that cells that express TLR4/MD-2, but not membrane-anchored CD14, will be sensitive to LPS but not S100A9.

The final implication is that sCD14 might have an important function in attenuating the activity of S100A9. Soluble CD14 circulates in the blood in healthy individuals at 2 mg/L (41–43). Because it inhibits the activation of TLR4 by S100A9, it could be part of mechanism to prevent over-activation of TLR4 by S100A9, thus preventing runaway inflammation. Taken together, these results suggest that CD14 may play an important role in regulating and tuning S100A9’s ability to activate TLR4.

### Interaction via a dynamic, multi-functional site

Putting together the complete mechanism by which S100A9 activates TLR4/MD-2 will require structural insight into how the protein promotes dimerization and, likely, internalization. We made a first step towards this goal by establishing how S100A9 interacts with CD14. S100A9 binds at a site that partially overlaps, but is distinct from, the LPS binding site on CD14. This region of CD14 is dynamic, fluctuating between an open and closed conformation (Figure 8D). The closed conformation is favored in the absence of ligand, shielding the hydrophobic pocket. The open conformation is favored when LPS binds because it sequesters the hydrophobic acyl chains of LPS away from water. Finally, both conformations are populated when S100A9 binds. The plastic nature of this region allows CD14 to bind to both LPS and S100A9, despite their low chemical similarity.

The complicated dynamics of this region may also explain our experimental results. Mutations of both Surface I and Surface II alter both activity and binding. This strongly implicates both surfaces in the S100A9/CD14 interaction. However, their effects were epistatic and thus difficult to rationalize in terms of simple charge (e.g., Arg to Glu) or size (e.g., Phe to Ala) reversals. If S100A9 does indeed interact with CD14 via two binding modes, mutations may perturb one mode and not the other. Or, more confusingly, a mutation that disrupts one binding mode might enhance another. This means a full dissection of the binding surface will likely require a combinatorial set of mutations to fully disrupt the network of interactions predicted by the MD simulations. We may also need to characterize multiple amino acid substitutions at each site, allowing us to probe the importance of specific physiochemical features at each site. Although we have yet to completely dissect this interface, we have identified a plausible route for the complete separation of function for LPS and S100A9.

### Future steps

The long-term goal of this project is to understand how S100A9 activates TLR4/MD-2. Our work provides a stepping-stone on the way to this outcome, as we can now ask how CD14 and S100A9, together, activate the complex. This provides a more nuanced starting point than just generically considering S100A9. For example, if the CD14/S100A9 complex physically interacts with TLR4/MD-2—as seems plausible—our work provides strong constraints on the orientation of the initial encounter, as we know where S100A9 interacts with CD14 and that both CD14 and TLR4/MD-2 are membrane anchored. This also sharpens our mechanistic questions. Rather than asking how S100A9 activates TLR4/MD-2, we can instead ask how CD14/S100A9 together promote internalization. Answering such questions will be critically important for understanding the basis for this biologically important, but poorly understood, proinflammatory mechanism.

## EXPERIMENTAL PROCEDURES

### Heterologous S100A9 expression and purification from E. coli

We expressed and purified S100A9 as previously described (14). Briefly, we expressed cysteine-free human S100A9 (C3S) in the pETDUET-1 vector. We transformed *E. coli* Rosetta BL21(DE3) *pLysS*. We grew 1.5 L liquid cultures to an OD_600_ ∼ 0.8 and then induced with 1 mM IPTG for 16 hours at 4°C. We harvested cells by centrifugation and lysed them via sonication. We purified S100A9 using three chromatography steps: immobilized metal ion affinity (HisTrap) at pH 7.4, anion exchange (HiTrap Q) at pH 8, followed by another anion exchange (HiTrap Q) at pH 6. We verified protein purity was >95% by SDS-PAGE. We concentrated and buffer exchanged proteins into 25 mM Tris, 100 mM NaCl, pH 7.4, then flash-froze dropwise into liquid nitrogen. Proteins were stored at −80 °C until needed. We determined protein concentration using A_280_ with an extinction coefficient of 6990 M^−1^ cm^−1^ (monomer). The protein concentrations reported in this manuscript are μM dimer.

### Heterologous sCD14 expression and purification from HEK293F cells

We expressed soluble CD14 and its mutants in Freestyle HEK293F cells. We started with the full-length human CD14 gene in the pcDNA3 backbone. pcDNA3-CD14 was a gift from Doug Golenbock (Addgene plasmid # 13645; http://n2t.net/addgene:13645; RRID:Addgene_13645). We designed primers to truncate CD14 at residue 367 (truncation at this residue to produce soluble CD14 has been previously reported (46)) and add a 6× His tag.

HEK293F cells were maintained in Freestyle293 Expression Medium, shaking at 135 rpm at 37 °C in 5% CO_2_. We transfected the HEK293F cells and expressed protein following the manufacturer’s instructions. Twenty-four hours prior to transfection, we seeded cells at 6-7 × 10^5^ cells/mL in Freestyle 293F Expression Medium. The cell viability at the time of transfection was >90% as determined by trypan blue staining. We transfected 30 mL of cells at 1 × 10^6^ cells/mL, using 293fectin and 1 μg of DNA per mL of cells. After 7 days of transfection, we harvested the supernatant and incubated it with 1 mL of Ni-NTA agarose at 4 °C for 1 hour. We then washed the resin twice with 25 mL of PBS containing 25 mM imidazole. We eluted protein with 5 mL PBS containing 500 mM imidazole. We buffer exchanged and concentrated the protein into PBS using Pall Microsep spin columns, then flash froze dropwise into liquid nitrogen. We measured the protein concentration by A_280_, using the calculated extinction coefficient of 31105 M^−1^ cm^−1^.

### NF-κB activity assay

We assayed NF-κB activity as previously described (13, 14, 37). We maintained HEK293T cells up to 30 passages in DMEM + 10% FBS + Antibiotic-Antimycotic, at 37 °C in 5% CO_2_. When performing an assay, we transiently transfected these cells with appropriate plasmids in a 96-well tissue culture plate using Lipofectamine and Plus (ThermoFisher). For the receptor complex, we used pcDNA3 plasmids that individually encoded full-length human TLR4, MD-2, and CD14 genes under control of the CMV constitutive promoter. hTLR4 was a gift from Ruslan Medzhitov (Addgene plasmid # 13086; http://n2t.net/addgene:13086; RRID:Addgene_13086). hMD-2_pcDNA3.1+ was purchased from Genscript (LY96_OHu26610C_pcDNA3.1(+)). pcDNA3-CD14 was a gift from Doug Golenbock (Addgene plasmid # 13645; http://n2t.net/addgene:13645; RRID:Addgene_13645). To measure NF-κB activity, we used the pGL3-elam-luc plasmid, which encodes the firefly luciferase behind one NF-κB promoter, and the pRL-TK plasmid, which encodes renilla luciferase behind the TK constitutive promoter. pGL3-ELAM-luc was a gift from Doug Golenbock (Addgene plasmid # 13029; http://n2t.net/addgene:13029; RRID:Addgene_13029). pRL-TK was purchased from Promega. We transfected a total of 100 ng DNA per well, diluted in Opti-Mem. The mix included 69 ng of empty pcDNA3 vector, as well as our experimental plasmids in the following amounts: 10 ng hTLR4, 0.5 ng hMD-2, 0.25 ng hCD14, 0.25 ng Renilla luciferase, and 20 ng firefly luciferase. We combined 65 μL of transfection mix with 135 μL of DMEM + FBS in each well. After 20 hours of transfection, we removed the transfection mix and replaced it with 100 μL/well of treatment.

Treatment mixes contained 75 μL DMEM and 25 μL treatment. We used *E. coli* K12 LPS (tlrl-klps, Invivogen). S100A9 was buffer exchanged into endotoxin-free PBS prior to treatment, using Pall Microsep concentrator spin columns. All S100A9 treatments included 200 μg/mL polymyxin B. We diluted treatment components to their desired concentration in endotoxin-free PBS (without Ca^2+^ or Mg^2+^). After three hours of treatment, we measured luciferase activity using the Dual-Glo Luciferase Assay Kit (Promega). All measurements were performed in technical triplicate. For data processing and normalization between experiments, each plate contained the following four treatments applied to the cells transfected with the complete human TLR4/MD-2/CD14 complex: mock (PBS), 200 ng/mL LPS, 200 ng/mL LPS with 200 μg/mL polymyxin B, and 2 μM S100A9 with 200 μg/mL polymyxin B.

After reading each plate, we took the mean of the technical replicates for each condition. To control for transfection efficiency, we normalized our firefly luciferase signal (FF) to our Renilla luciferase signal. For a given sample and treatment *i*, the activity was:

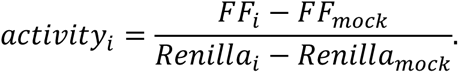

To allow comparison of activity between plates collected on different days, we additionally normalized the activity of each sample and treatment to the wildtype TLR4/MD-2/CD14 response 200 ng/mL LPS or 2 μM S100A9:

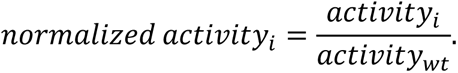

All plots and analysis report the normalized activity.

### Docking with AlphaFold2

We used the AlphaFold2 Google Colab notebook to predict the structure of the CD14/S100A9 complex (53–55). We input a single CD14 sequence (UniProt ID: P08571) and two S100A9 sequences (Uniprot ID: P06702), thus setting the expected stoichiometry to one CD14 interacting with an S100A9 dimer. We used the top-ranked model for all further analyses.

### BioLayer Interferometry (Gator)

We purified S100A9/C3S/P114C expressed in *E. coli* using our standard method, then biotinylated the protein using the EZ-link BMCC kit, following the manufacturer’s instructions. We confirmed biotinylation by MALDI-TOF mass spectrometry. The binding assays were performed using the GatorPrime BLI. Our running buffer consisted of 25 mM Tris (pH 7.4), 100 mM NaCl, 2 mM CaCl_2_, 1% BSA, and 0.1% Tween-20. We included BSA and Tween-20 to block non-specific binding. All proteins were buffer exchanged into running buffer prior to our experiments. We soaked streptavidin sensors in running buffer for a minimum of 15 minutes prior to our experiments to dissolve sucrose coating. For each experiment, we sequentially dipped sensors into wells of a 96-well plate containing the following conditions for the indicated times: buffer for 30 s, S100A9 at 50 nM for 120 s, buffer for 30 s, sCD14 at 0.5, 1.0, 2.0, or 4.0 μM for 240s, then buffer for 240 s. We ran our experiments at 30 °C, shaking at 1000 rpm during reads. Binding fitting was performed with the GatorBio Software.

### Model construction for molecular dynamics simulations

For the CD14-alone simulations, we started with the crystal structure of the human protein (RCSB ID: 4GLP, (25)). In the crystal structure, there is a deep hydrophobic pocket exposed to solvent on the N-terminus of the protein. Within 5 ns of simulation time, the helix formed by residues 26-55 closed over the pocket, expelling water molecules and burying the hydrophobic surface of the pocket (Figure S2). This can be seen in Figure 6A, which shows CD14 in the closed conformation. Residues 26-55 (yellow) act as a “lid” that fills the pocket.

For the LPS simulations, we studied the interaction of CD14 with *E. coli* Lipid A (LA). LA has the six acyl chains, three glucosamines, and two phosphates of *E. coli* LPS, but does not possess the core sugars or O-antigen. Lipid A is the conserved molecular pattern recognized by TLR4/MD-2/CD14 (58). We did not include the core or O-antigen because they consist of long, dynamic polymers of sugar moieties that would have been computationally expensive to model. To dock LA into CD14, we oriented LA with its acyl chains facing the CD14 binding pocket but with no atom closer than 5 Å. We then ran short (20 ns) docking simulations. We started ten simulations with the crystal structure of CD14 and another ten with a CD14 structure pre-equilibrated by 100 ns of MD simulation. We found that LA engaged with and inserted into the CD14 pocket in 9 out of 10 of the crystal structure simulations and in 3 of the 10 pre-equilibrated simulations. This difference in success rate is because the lid residues started in the open conformation in the crystal structure, but started closed in the pre-equilibrated structure. For production runs, we arbitrarily selected two of the LA docking runs that started from the crystal structure and one that started from the pre-equilibrated structure.

For the S100A9 simulations, we started with the AlphaFold structure (Figure 5A). We also started simulations with S100A9 and CD14 in a variety of different orientations relative to one another. This included three different models generated using RosettaDock (59), three models with S100A9 in the same orientation as in the AlphaFold structure, but with CD14 in the closed state (achieved by pre-equilibration with 100 ns of simulation), and three models with S100A9 in an arbitrary orientation relative to the CD14 crystal structure. Only simulations that started with the AlphaFold CD14/S100A9 model were stable; in all other simulations, the S100A9 and CD14 drifted apart within ∼50 ns. We therefore performed our CD14/S100A9 simulations using the AlphaFold structure as our starting conformation.

### Molecular dynamics simulation parameters

For all simulations, we used GROMACS 2023 (60, 61) with the CHARMM36 2021 forcefield (52) and TIP3P waters (62). We generated lipid A coordinates and forcefield parameters using LPS Modeler (63) as implemented in CHARMM-GUI (64). We placed Ca^2+^ ions in the AlphaFold structure of S100A9 by aligning the crystal structure of Ca^2+^-bound S100A9 (RCSB ID: 1IRJ (33)) and then manually extracting the Ca^2+^ coordinates. Using the GROMACS pdb2gmx module, we constructed a cubic periodic solvent box 20 Å longer than the maximum model dimension, neutralized the system by randomly placing 100 mM Na^+^/Cl^−^ counter ions, added missing hydrogen atoms, and assigned protonation states at a pH of 7.0. We prepared the system for simulations with three final steps: 1) Steepest descent energy minimization; 2) 100 ps of position-restrained equilibration in the NVT ensemble (assigning initial velocities from a Maxwell distribution at 300 K); and 3) 100 ps of equilibration in the NPT ensemble. We restrained the positions of all non-solvent heavy atoms in our position-restrained simulations. We did our production runs using an NPT ensemble at 1 atmosphere and 300 K. We used isotropic Parrinello-Rahman pressure coupling (65, 66) and velocity-rescaling temperature coupling (67). We used LINCS for bond constraints (68, 69), treated non-bonded interactions with a Verlet scheme (70), and captured long-range electrostatics using a 4^th^-order Particle Mesh Ewald approximation (71). We did all calculations using the talapas high-performance computing cluster at the University of Oregon

### Analysis of MD trajectories

We analyzed the results using a Visual Molecular Dynamics (72), python scripts using the MDAnalysis library (73, 74), and PyMOL (75). We calculated solvent-accessible surface areas using the freesasa library (76) with a solvent radius of 1.4 Å.

For our clustering analysis (Figure 7), we went through our simulations, calculating contacts, hydrogen bonds, and the orientation of S100A9 relative to CD14 for ∼270,000 frames. We defined two residues as in contact if they that had at least one pair of non-hydrogen atoms within 4 Å. We measured hydrogen bonds using the MDAnalysis HydrogenBondAnalysis package (77), defining hydrogen bonds as an oxygen or nitrogen within 3.0 Å of a polar proton where the donor/proton/acceptor angle was at least 150°. We calculated the orientation of S100A9 relative to CD14 in two steps. First, we aligned each frame in the simulation to the starting frame using the C_α_ atoms of CD14 residues 100-200. This central region of CD14 is rigid and thus provides an approximately fixed reference against which to measure the orientation of S100A9. We then calculated the rotation matrix that would minimize the root-mean squared deviation of S100A9 from the CD14-aligned frame to S100A9 from the initial conformation (78, 79). We limited this analysis to the C_α_ atoms from residues 4-90 from both chains, as these residues were ordered throughout the simulations. The resulting rotation matrix has nine values; we treated each as its own feature in the downstream analysis.

We then encoded contacts, hydrogen bonds, and orientation of S100A9 as features. We scored each contact and hydrogen bond as present (1) or absent (0) in each frame. The raw orientation of S100A9 in each frame was given by nine floating point numbers between −1 and 1. We rescaled these values to be between 0-1 to match the scale of the hydrogen bond and contact features. We then applied a 100-frame sliding window. This transformed our Boolean contact and hydrogen bond scores to float values between 0 and 1. We then identified the set of hydrogen bonds and contacts that accounted for the top 99% of observed contact density. We dropped any hydrogen bonds or contacts outside this set. This yielded 228 final features: 86 hydrogen bonds, 133 contacts, and 9 rotation matrix elements. Finally, we sampled these features every 100^th^ frame—matching the size of our smoothing window—yielding ∼2,700 frames, each with 228 features.

We calculated the Euclidean distance between all frames and clustered using k-means as implemented in scikit-learn (80). We generated between 2 and 20 clusters and selected the cluster with the highest Silhouette Score (81). This proved to be two clusters (score of 0.35). We checked for robustness to feature choice by re-doing the analysis excluding all hydrogen bonds, contacts, or orientation scores. We also used k-fold cross validation (k=10) sampling from all 228 features. In every case, we found two clusters with similar structural features in similar proportions across the simulations.

### Table of key reagents used in this study

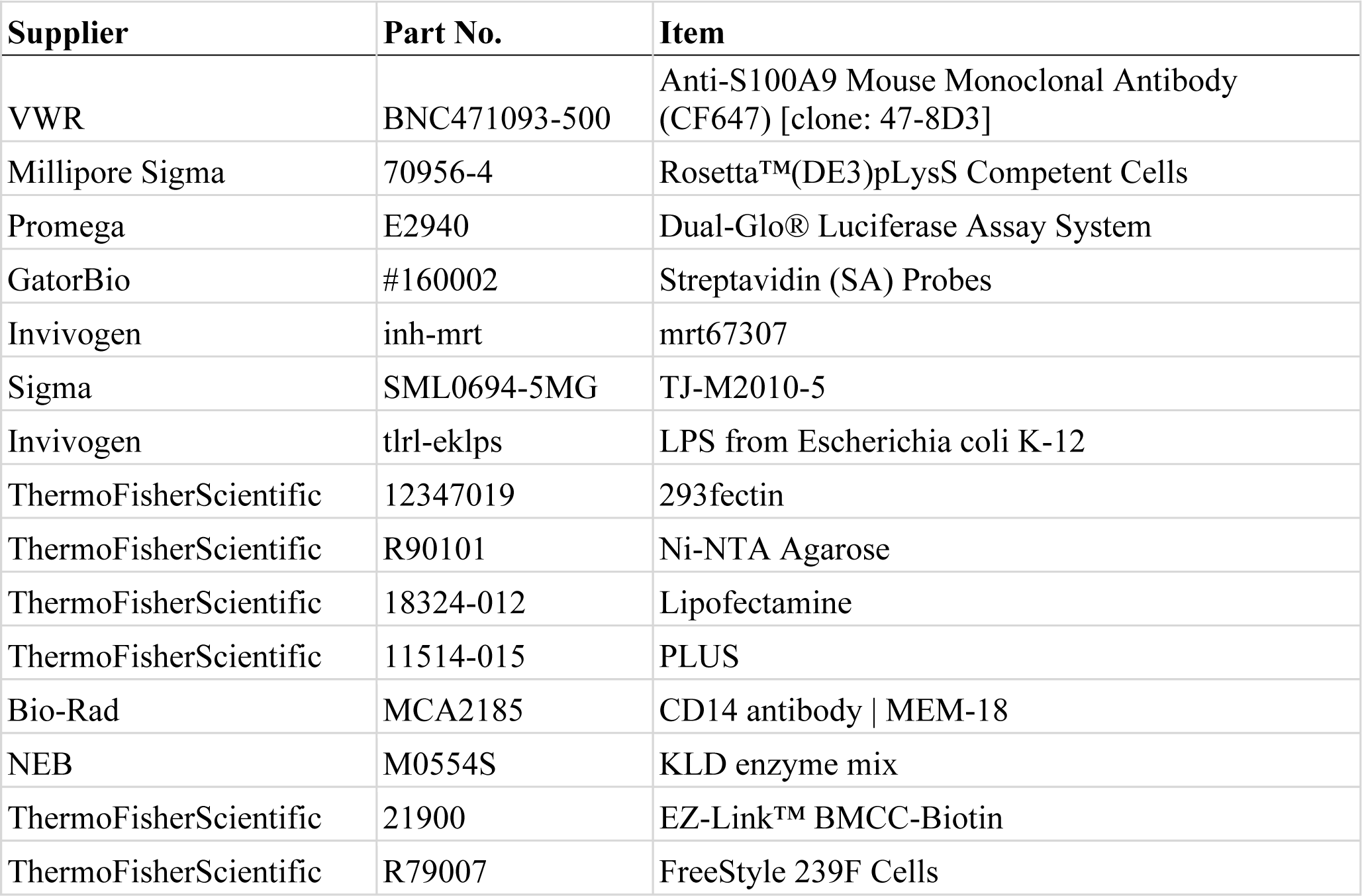

| **Supplier** | **Part No.** | **Item** |
| --- | --- | --- |
| VWR | BNC471093-500 | Anti-S100A9 Mouse Monoclonal Antibody (CF647) [clone: 47-8D3] |
| Millipore Sigma | 70956-4 | Rosetta™(DE3)pLysS Competent Cells |
| Promega | E2940 | Dual-Glo® Luciferase Assay System |
| GatorBio | #160002 | Streptavidin (SA) Probes |
| Invivogen | inh-mrt | mrt67307 |
| Sigma | SML0694-5MG | TJ-M2010-5 |
| Invivogen | tlrl-eklps | LPS from Escherichia coli K-12 |
| ThermoFisherScientific | 12347019 | 293fectin |
| ThermoFisherScientific | R90101 | Ni-NTA Agarose |
| ThermoFisherScientific | 18324-012 | Lipofectamine |
| ThermoFisherScientific | 11514-015 | PLUS |
| Bio-Rad | MCA2185 | CD14 antibody \| MEM-18 |
| NEB | M0554S | KLD enzyme mix |
| ThermoFisherScientific | 21900 | EZ-Link™ BMCC-Biotin |
| ThermoFisherScientific | R79007 | FreeStyle 239F Cells |

## Data Availability Supplement

All data described in the manuscript are included in the figures and the supplemental materials.

## Acknowledgements

We thank current and former members of the Harms lab for helpful discussion and input. We thank the Barber lab for use of their cell culture facility and gift of HEK293F cells. This work benefited from access to the University of Oregon high performance computing cluster, Talapas. Funding: 5T32GM007759 (LOC), The John Keana Graduate Student Fellowship (LOC), and NIGMS R01-GM146114 (MJH).

## References

1. Berntzen, H. B., and Fagerhol, M. K. (1990) L1, a major granulocyte protein; isolation of high quantities of its subunits. Scandinavian Journal of Clinical and Laboratory Investigation. 50, 769–774

2. Chen, B., Miller, A. L., Rebelatto, M., Brewah, Y., Rowe, D. C., Clarke, L., Czapiga, M., Rosenthal, K., Imamichi, T., Chen, Y., Chang, C.-S., Chowdhury, P. S., Naiman, B., Wang, Y., Yang, D., Humbles, A. A., Herbst, R., and Sims, G. P. (2015) S100A9 Induced Inflammatory Responses Are Mediated by Distinct Damage Associated Molecular Patterns (DAMP) Receptors In Vitro and In Vivo. PLOS ONE. 10, e0115828

3. Ryckman, C., Vandal, K., Rouleau, P., Talbot, M., and Tessier, P. A. (2003) Proinflammatory activities of S100: proteins S100A8, S100A9, and S100A8/A9 induce neutrophil chemotaxis and adhesion. J Immunol. 170, 3233–3242

4. Averill Michelle M., Kerkhoff Claus, and Bornfeldt Karin E. (2012) S100A8 and S100A9 in Cardiovascular Biology and Disease. Arteriosclerosis, Thrombosis, and Vascular Biology. 32, 223–229

5. Markowitz, J., and Carson, W. E. (2013) Review of S100A9 biology and its role in cancer. Biochimica et Biophysica Acta (BBA) - Reviews on Cancer. 1835, 100–109

6. Ehrchen, J. M., Sunderkötter, C., Foell, D., Vogl, T., and Roth, J. (2009) The endogenous Toll–like receptor 4 agonist S100A8/S100A9 (calprotectin) as innate amplifier of infection, autoimmunity, and cancer. Journal of Leukocyte Biology. 86, 557–566

7. Hermani, A., Hess, J., Servi, B. D., Medunjanin, S., Grobholz, R., Trojan, L., Angel, P., and Mayer, D. (2005) Calcium-Binding Proteins S100A8 and S100A9 as Novel Diagnostic Markers in Human Prostate Cancer. Clin Cancer Res. 11, 5146–5152

8. 8. Zhang, C., Liu, Y., Gilthorpe, J., and van der Maarel, J. R. C. (2012) MRP14 (S100A9) Protein Interacts with Alzheimer Beta-Amyloid Peptide and Induces Its Fibrillization. PLoS One. 10.1371/journal.pone.0032953

9. Wang, C., Iashchishyn, I. A., Pansieri, J., Nyström, S., Klementieva, O., Kara, J., Horvath, I., Moskalenko, R., Rofougaran, R., Gouras, G., Kovacs, G. G., Shankar, S. K., and Morozova-Roche, L. A. (2018) S100A9-Driven Amyloid-Neuroinflammatory Cascade in Traumatic Brain Injury as a Precursor State for Alzheimer’s Disease. Scientific Reports. 8, 12836

10. Zhang, X., Wei, L., Wang, J., Qin, Z., Wang, J., Lu, Y., Zheng, X., Peng, Q., Ye, Q., Ai, F., Liu, P., Wang, S., Li, G., Shen, S., and Ma, J. (2017) Suppression Colitis and Colitis-Associated Colon Cancer by Anti-S100a9 Antibody in Mice. Front Immunol. 8, 1774

11. Björk, P., Björk, A., Vogl, T., Stenström, M., Liberg, D., Olsson, A., Roth, J., Ivars, F., and Leanderson, T. (2009) Identification of Human S100A9 as a Novel Target for Treatment of Autoimmune Disease via Binding to Quinoline-3-Carboxamides. PLoS Biol. 10.1371/journal.pbio.1000097

12. Riva, M., Källberg, E., Björk, P., Hancz, D., Vogl, T., Roth, J., Ivars, F., and Leanderson, T. (2012) Induction of nuclear factor-κB responses by the S100A9 protein is Toll-like receptor-4-dependent. Immunology. 137, 172–182

13. Harman, J. L., Reardon, P. N., Costello, S. M., Warren, G. D., Phillips, S. R., Connor, P. J., Marqusee, S., and Harms, M. J. (2022) Evolution avoids a pathological stabilizing interaction in the immune protein S100A9. Proceedings of the National Academy of Sciences. 119, e2208029119

14. Harman, J. L., Loes, A. N., Warren, G. D., Heaphy, M. C., Lampi, K. J., and Harms, M. J. (2020) Evolution of multifunctionality through a pleiotropic substitution in the innate immune protein S100A9. eLife. 9, e54100

15. Pelletier, M., Simard, J.-C., Girard, D., and Tessier, P. A. (2018) Quinoline-3-carboxamides such as tasquinimod are not specific inhibitors of S100A9. Blood Adv. 2, 1170–1171

16. Raetz, C. R. H., and Whitfield, C. (2002) Lipopolysaccharide Endotoxins. Annu Rev Biochem. 71, 635–700

17. Galloway, S. M., and Raetz, C. R. (1990) A mutant of Escherichia coli defective in the first step of endotoxin biosynthesis. Journal of Biological Chemistry. 265, 6394–6402

18. Nagai, Y., Akashi, S., Nagafuku, M., Ogata, M., Iwakura, Y., Akira, S., Kitamura, T., Kosugi, A., Kimoto, M., and Miyake, K. (2002) Essential role of MD-2 in LPS responsiveness and TLR4 distribution. Nat Immunol. 3, 667–672

19. da Silva Correia, J., Soldau, K., Christen, U., Tobias, P. S., and Ulevitch, R. J. (2001) Lipopolysaccharide is in close proximity to each of the proteins in its membrane receptor complex. transfer from CD14 to TLR4 and MD-2. J Biol Chem. 276, 21129–21135

20. Jagtap, P., Prasad, P., Pateria, A., Deshmukh, S. D., and Gupta, S. (2020) A Single Step in vitro Bioassay Mimicking TLR4-LPS Pathway and the Role of MD2 and CD14 Coreceptors. Front Immunol. 10.3389/fimmu.2020.00005

21. Huber, R. G., Berglund, N. A., Kargas, V., Marzinek, J. K., Holdbrook, D. A., Khalid, S., Piggot, T. J., Schmidtchen, A., and Bond, P. J. (2018) A Thermodynamic Funnel Drives Bacterial Lipopolysaccharide Transfer in the TLR4 Pathway. Structure. 26, 1151–1161.e4

22. Wright, S. D., Ramos, R. A., Tobias, P. S., Ulevitch, R. J., and Mathison, J. C. (1990) CD14, a Receptor for Complexes of Lipopolysaccharide (LPS) and LPS Binding Protein. Science. 249, 1431–1433

23. Cunningham, M. D., Shapiro, R. A., Seachord, C., Ratcliffe, K., Cassiano, L., and Darveau, R. P. (2000) CD14 Employs Hydrophilic Regions to “Capture” Lipopolysaccharides. The Journal of Immunology. 164, 3255–3263

24. Kim, J.-I., Lee, C. J., Jin, M. S., Lee, C.-H., Paik, S.-G., Lee, H., and Lee, J.-O. (2005) Crystal Structure of CD14 and Its Implications for Lipopolysaccharide Signaling. J. Biol. Chem. 280, 11347–11351

25. Kelley, S. L., Lukk, T., Nair, S. K., and Tapping, R. I. (2013) The Crystal Structure of Human Soluble CD14 Reveals a Bent Solenoid with a Hydrophobic Amino-Terminal Pocket. The Journal of Immunology. 190, 1304–1311

26. Ryu, J.-K., Kim, S. J., Rah, S.-H., Kang, J. I., Jung, H. E., Lee, D., Lee, H. K., Lee, J.-O., Park, B. S., Yoon, T.-Y., and Kim, H. M. (2017) Reconstruction of LPS Transfer Cascade Reveals Structural Determinants within LBP, CD14, and TLR4-MD2 for Efficient LPS Recognition and Transfer. Immunity. 46, 38–50

27. Hailman, E., Lichenstein, H. S., Wurfel, M. M., Miller, D. S., Johnson, D. A., Kelley, M., Busse, L. A., Zukowski, M. M., and Wright, S. D. (1994) Lipopolysaccharide (LPS)-binding protein accelerates the binding of LPS to CD14. Journal of Experimental Medicine. 179, 269–277

28. Haziot, A., Chen, S., Ferrero, E., Low, M. G., Silber, R., and Goyert, S. M. (1988) The monocyte differentiation antigen, CD14, is anchored to the cell membrane by a phosphatidylinositol linkage. J Immunol. 141, 547–552

29. Simmons, D., Tan, S., Tenen, D., Nicholson-Weller, A., and Seed, B. (1989) Monocyte antigen CD14 is a phospholipid anchored membrane protein. Blood. 73, 284–289

30. Zanoni, I., Ostuni, R., Marek, L. R., Barresi, S., Barbalat, R., Barton, G. M., Granucci, F., and Kagan, J. C. (2011) CD14 Controls the LPS-Induced Endocytosis of Toll-like Receptor 4. Cell. 147, 868–880

31. Ciesielska, A., Matyjek, M., and Kwiatkowska, K. (2021) TLR4 and CD14 trafficking and its influence on LPS-induced pro-inflammatory signaling. Cell Mol Life Sci. 78, 1233–1261

32. Jiang, Z., Georgel, P., Du, X., Shamel, L., Sovath, S., Mudd, S., Huber, M., Kalis, C., Keck, S., Galanos, C., Freudenberg, M., and Beutler, B. (2005) CD14 is required for MyD88-independent LPS signaling. Nat Immunol. 6, 565–570

33. Itou, H., Yao, M., Fujita, I., Watanabe, N., Suzuki, M., Nishihira, J., and Tanaka, I. (2002) The crystal structure of human MRP14 (S100A9), a Ca2+-dependent regulator protein in inflammatory process11Edited by R. Huber. Journal of Molecular Biology. 316, 265–276

34. Park, B. S., Song, D. H., Kim, H. M., Choi, B.-S., Lee, H., and Lee, J.-O. (2009) The structural basis of lipopolysaccharide recognition by the TLR4-MD-2 complex. Nature. 458, 1191–1195

35. PubChem GPI-anchor amidated glycine. [online] https://pubchem.ncbi.nlm.nih.gov/compound/145996624 (Accessed April 1, 2024)

36. He, Z., Riva, M., Björk, P., Swärd, K., Mörgelin, M., Leanderson, T., and Ivars, F. (2016) CD14 Is a Co-Receptor for TLR4 in the S100A9-Induced Pro-Inflammatory Response in Monocytes. PLoS One. 10.1371/journal.pone.0156377

37. Loes, A. N., Bridgham, J. T., and Harms, M. J. (2018) Coevolution of the Toll-Like Receptor 4 Complex with Calgranulins and Lipopolysaccharide. Front Immunol. 10.3389/fimmu.2018.00304

38. Anderson, J. A., Loes, A. N., Waddell, G. L., and Harms, M. J. (2019) Tracing the evolution of novel features of human Toll-like receptor 4. Protein Science. 28, 1350–1358

39. Rouillard, A. D., Gundersen, G. W., Fernandez, N. F., Wang, Z., Monteiro, C. D., McDermott, M. G., and Ma’ayan, A. (2016) The harmonizome: a collection of processed datasets gathered to serve and mine knowledge about genes and proteins. Database. 2016, baw100

40. Yu, B., and Wright, S. D. (1996) Catalytic properties of lipopolysaccharide (LPS) binding protein. Transfer of LPS to soluble CD14. J Biol Chem. 271, 4100–4105

41. Lien, E., Aukrust, P., Sundan, A., Müller, F., Frøland, S. S., and Espevik, T. (1998) Elevated levels of serum-soluble CD14 in human immunodeficiency virus type 1 (HIV-1) infection: correlation to disease progression and clinical events. Blood. 92, 2084–2092

42. Sandler, N. G., Wand, H., Roque, A., Law, M., Nason, M. C., Nixon, D. E., Pedersen, C., Ruxrungtham, K., Lewin, S. R., Emery, S., Neaton, J. D., Brenchley, J. M., Deeks, S. G., Sereti, I., and Douek, D. C. (2011) Plasma Levels of Soluble CD14 Independently Predict Mortality in HIV Infection. J Infect Dis. 203, 780–790

43. Gonzalez-Quintela, A., Alonso, M., Campos, J., Vizcaino, L., Loidi, L., and Gude, F. (2013) Determinants of serum concentrations of lipopolysaccharide-binding protein (LBP) in the adult population: the role of obesity. PLoS One. 8, e54600

44. Mukherjee, R., Kanti Barman, P., Kumar Thatoi, P., Tripathy, R., Kumar Das, B., and Ravindran, B. (2015) Non-Classical monocytes display inflammatory features: Validation in Sepsis and Systemic Lupus Erythematous. Sci Rep. 5, 13886

45. Frey, E. A., Miller, D. S., Jahr, T. G., Sundan, A., Bazil, V., Espevik, T., Finlay, B. B., and Wright, S. D. (1992) Soluble CD14 participates in the response of cells to lipopolysaccharide. Journal of Experimental Medicine. 176, 1665–1671

46. Juan, T. S.-C., Kelley, M. J., Johnson, D. A., Busse, L. A., Hailman, E., Wright, S. D., and Lichenstein, H. S. (1995) Soluble CD14 Truncated at Amino Acid 152 Binds Lipopolysaccharide (LPS) and Enables Cellular Response to LPS (∗). Journal of Biological Chemistry. 270, 1382–1387

47. Kawai, T., and Akira, S. (2010) The role of pattern-recognition receptors in innate immunity: update on Toll-like receptors. Nat Immunol. 11, 373–384

48. Petherick, K. J., Conway, O. J. L., Mpamhanga, C., Osborne, S. A., Kamal, A., Saxty, B., and Ganley, I. G. (2015) Pharmacological Inhibition of ULK1 Kinase Blocks Mammalian Target of Rapamycin (mTOR)-dependent Autophagy. J Biol Chem. 290, 11376–11383

49. Clark, K., Peggie, M., Plater, L., Sorcek, R. J., Young, E. R. R., Madwed, J. B., Hough, J., McIver, E. G., and Cohen, P. (2011) Novel cross-talk within the IKK family controls innate immunity. Biochem J. 434, 93–104

50. Xie, L., Jiang, F.-C., Zhang, L.-M., He, W.-T., Liu, J.-H., Li, M.-Q., Zhang, X., Xing, S., Guo, H., and Zhou, P. (2016) Targeting of MyD88 Homodimerization by Novel Synthetic Inhibitor TJ-M2010-5 in Preventing Colitis-Associated Colorectal Cancer. JNCI: Journal of the National Cancer Institute. 108, djv364

51. Miao, Y., Ding, Z., Zou, Z., Yang, Y., Yang, M., Zhang, X., Li, Z., Zhou, L., Zhang, L., Zhang, X., Du, D., Jiang, F., and Zhou, P. (2020) Inhibition of MyD88 by a novel inhibitor reverses two-thirds of the infarct area in myocardial ischemia and reperfusion injury. Am J Transl Res. 12, 5151–5169

52. Juan, T. S.-C., Hailman, E., Kelley, M. J., Busse, L. A., Davy, E., Empig, C. J., Narhi, L. O., Wright, S. D., and Lichenstein, H. S. (1995) Identification of a Lipopolysaccharide Binding Domain in CD14 between Amino Acids 57 and 64 (∗). Journal of Biological Chemistry. 270, 5219–5224

53. Mirdita, M., Schütze, K., Moriwaki, Y., Heo, L., Ovchinnikov, S., and Steinegger, M. (2022) ColabFold: making protein folding accessible to all. Nat Methods. 19, 679–682

54. Evans, R., O’Neill, M., Pritzel, A., Antropova, N., Senior, A., Green, T., Žídek, A., Bates, R., Blackwell, S., Yim, J., Ronneberger, O., Bodenstein, S., Zielinski, M., Bridgland, A., Potapenko, A., Cowie, A., Tunyasuvunakool, K., Jain, R., Clancy, E., Kohli, P., Jumper, J., and Hassabis, D. (2022) Protein complex prediction with AlphaFold-Multimer. 10.1101/2021.10.04.463034

55. Jumper, J., Evans, R., Pritzel, A., Green, T., Figurnov, M., Ronneberger, O., Tunyasuvunakool, K., Bates, R., Žídek, A., Potapenko, A., Bridgland, A., Meyer, C., Kohl, S. A. A., Ballard, A. J., Cowie, A., Romera-Paredes, B., Nikolov, S., Jain, R., Adler, J., Back, T., Petersen, S., Reiman, D., Clancy, E., Zielinski, M., Steinegger, M., Pacholska, M., Berghammer, T., Bodenstein, S., Silver, D., Vinyals, O., Senior, A. W., Kavukcuoglu, K., Kohli, P., and Hassabis, D. (2021) Highly accurate protein structure prediction with AlphaFold. Nature. 596, 583–589

56. Bonhomme, D., Santecchia, I., Vernel-Pauillac, F., Caroff, M., Germon, P., Murray, G., Adler, B., Boneca, I. G., and Werts, C. (2020) Leptospiral LPS escapes mouse TLR4 internalization and TRIF-associated antimicrobial responses through O antigen and associated lipoproteins. PLoS Pathog. 10.1371/journal.ppat.1008639

57. Tan, Y., Zanoni, I., Cullen, T. W., Goodman, A. L., and Kagan, J. C. (2015) Mechanisms of Toll-like receptor 4 endocytosis reveal a common immune-evasion strategy used by pathogenic and commensal bacteria. Immunity. 43, 909–922

58. Bryant, C. E., Spring, D. R., Gangloff, M., and Gay, N. J. (2010) The molecular basis of the host response to lipopolysaccharide. Nat Rev Microbiol. 8, 8–14

59. Wang, C., Bradley, P., and Baker, D. (2007) Protein-protein docking with backbone flexibility. J Mol Biol. 373, 503–519

60. Abraham, M. J., Murtola, T., Schulz, R., Páll, S., Smith, J. C., Hess, B., and Lindahl, E. (2015) GROMACS: High performance molecular simulations through multi-level parallelism from laptops to supercomputers. SoftwareX. 1–2, 19–25

61. Van Der Spoel, D., Lindahl, E., Hess, B., Groenhof, G., Mark, A. E., and Berendsen, H. J. C. (2005) GROMACS: Fast, flexible, and free. Journal of Computational Chemistry. 26, 1701–1718

62. MacKerell, A. D., Bashford, D., Bellott, M., Dunbrack, R. L., Evanseck, J. D., Field, M. J., Fischer, S., Gao, J., Guo, H., Ha, S., Joseph-McCarthy, D., Kuchnir, L., Kuczera, K., Lau, F. T., Mattos, C., Michnick, S., Ngo, T., Nguyen, D. T., Prodhom, B., Reiher, W. E., Roux, B., Schlenkrich, M., Smith, J. C., Stote, R., Straub, J., Watanabe, M., Wiórkiewicz-Kuczera, J., Yin, D., and Karplus, M. (1998) All-atom empirical potential for molecular modeling and dynamics studies of proteins. J Phys Chem B. 102, 3586–3616

63. Lee, J., Patel, D. S., Ståhle, J., Park, S.-J., Kern, N. R., Kim, S., Lee, J., Cheng, X., Valvano, M. A., Holst, O., Knirel, Y. A., Qi, Y., Jo, S., Klauda, J. B., Widmalm, G., and Im, W. (2019) CHARMM-GUI Membrane Builder for Complex Biological Membrane Simulations with Glycolipids and Lipoglycans. J. Chem. Theory Comput. 15, 775–786

64. Jo, S., Kim, T., Iyer, V. G., and Im, W. (2008) CHARMM-GUI: A web-based graphical user interface for CHARMM. Journal of Computational Chemistry. 29, 1859–1865

65. Parrinello, M., and Rahman, A. (1982) Strain fluctuations and elastic constants. The Journal of Chemical Physics. 76, 2662–2666

66. Nosé, S., and Klein, M. L. (1983) Constant pressure molecular dynamics for molecular systems. Molecular Physics. 50, 1055–1076

67. Bussi, G., Donadio, D., and Parrinello, M. (2007) Canonical sampling through velocity rescaling. The Journal of Chemical Physics. 126, 014101

68. Hess, B., Bekker, H., Berendsen, H. J. C., and Fraaije, J. G. E. M. (1997) LINCS: A linear constraint solver for molecular simulations. Journal of Computational Chemistry. 18, 1463–1472

69. Hess, B. (2008) P-LINCS: A Parallel Linear Constraint Solver for Molecular Simulation. J. Chem. Theory Comput. 4, 116–122

70. Swope, W. C., Andersen, H. C., Berens, P. H., and Wilson, K. R. (1982) A computer simulation method for the calculation of equilibrium constants for the formation of physical clusters of molecules: Application to small water clusters. The Journal of Chemical Physics. 76, 637–649

71. Darden, T., York, D., and Pedersen, L. (1993) Particle mesh Ewald: An N⋅log(N) method for Ewald sums in large systems. The Journal of Chemical Physics. 98, 10089–10092

72. Humphrey, W., Dalke, A., and Schulten, K. (1996) VMD: Visual molecular dynamics. Journal of Molecular Graphics. 14, 33–38

73. Michaud-Agrawal, N., Denning, E. J., Woolf, T. B., and Beckstein, O. (2011) MDAnalysis: A Toolkit for the Analysis of Molecular Dynamics Simulations. J Comput Chem. 32, 2319–2327

74. Gowers, R., Linke, M., Barnoud, J., Reddy, T., Melo, M., Seyler, S. L., Domański, J., Dotson, D., Buchoux, S., Kenney, I., and Beckstein, O. (2016) MDAnalysis: A Python Package for the Rapid Analysis of Molecular Dynamics Simulations, pp. 98–105, 10.25080/Majora-629e541a-00e

75. Schrödinger, LLC (2015) The PyMOL Molecular Graphics System, Version 1.8

76. Mitternacht, S. (2016) FreeSASA: An open source C library for solvent accessible surface area calculations. 10.12688/f1000research.7931.1

77. Smith, P., Ziolek, R. M., Gazzarrini, E., Owen, D. M., and Lorenz, C. D. (2019) On the interaction of hyaluronic acid with synovial fluid lipid membranes. Phys Chem Chem Phys. 21, 9845–9857

78. Theobald, D. L. (2005) Rapid calculation of RMSDs using a quaternion-based characteristic polynomial. Acta Crystallogr A. 61, 478–480

79. Liu, P., Agrafiotis, D. K., and Theobald, D. L. (2010) Fast determination of the optimal rotational matrix for macromolecular superpositions. J Comput Chem. 31, 1561–1563

80. Pedregosa, F., Varoquaux, G., Gramfort, A., Michel, V., Thirion, B., Grisel, O., Blondel, M., Prettenhofer, P., Weiss, R., Dubourg, V., Vanderplas, J., Passos, A., Cournapeau, D., Brucher, M., Perrot, M., and Duchesnay, É. (2011) Scikit-learn: Machine Learning in Python. Journal of Machine Learning Research. 12, 2825–2830

81. Rousseeuw, P. J. (1987) Silhouettes: A graphical aid to the interpretation and validation of cluster analysis. Journal of Computational and Applied Mathematics. 20, 53–65

