## Supplement for "S100A9 interacts with a dynamic region on CD14 to activate Toll-like receptor 4"

**Table S1: Effect of CD14 mutations on S100A9 and LPS activity.**

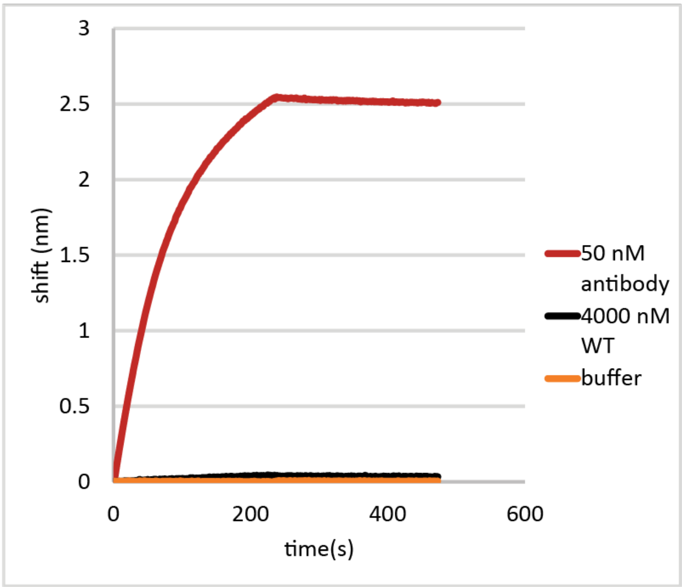


**Figure S1: mAB antibody reveals S100A9 was successfully on the BLI sensor.** Bio-Layer Interferometry binding data for immobilized S100A9 with different molecules flowed over the cell: 50 nM anti-S100A9 mAb (red), 4 µM sCD14 (black), and running buffer (orange). The S100A9 concentration was kept constant at 100 nM monomer. Binding plots are representative plots from one replicate. Binding was measured in triplicate at 50 nM and 100 nM mAb.

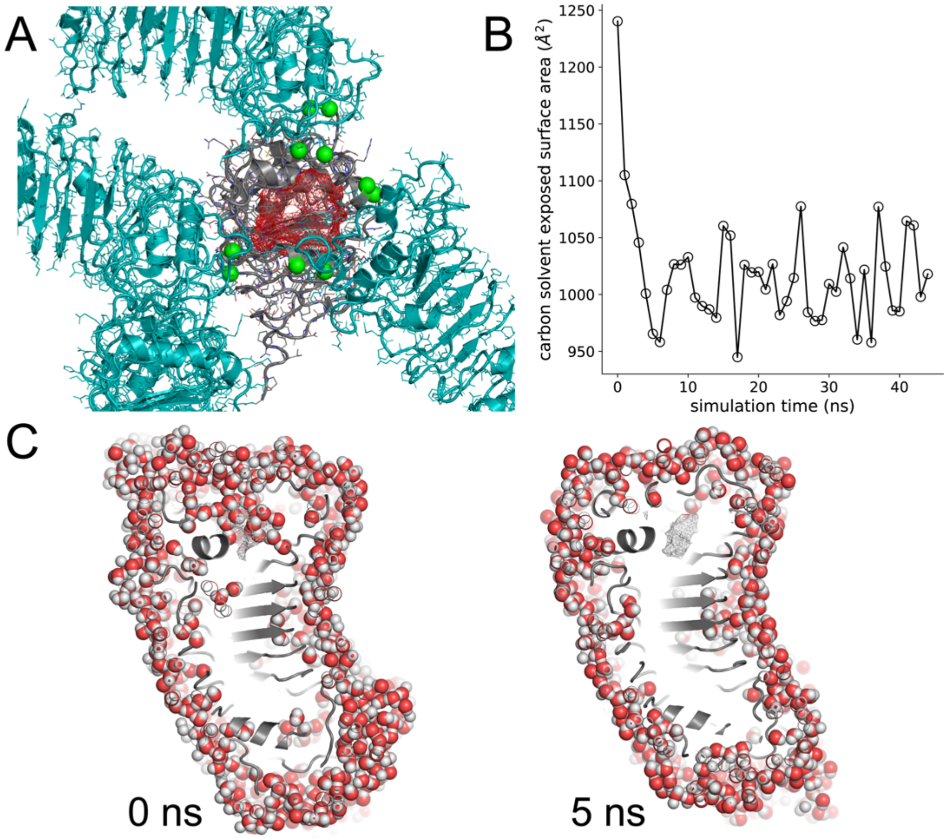


**Figure S2. The CD14 crystal structure has a dynamic hydrophobic pocket propped open by crystal contacts.** A) The crystal structure of human CD14 (RCSB ID: 4GLP). The CD14 monomer is the asymmetric unit, shown in gray. The four crystallographic symmetry mates adjacent to the binding pocket of the monomer are shown in teal. The green spheres are crystal contacts. In the crystal, CD14 has a large, concave hydrophobic surface (red mesh) surrounded by loops and helices supported by the crystal contacts. B) Plot shows solvent accessible surface area of CD14 carbons for snapshots taken ever nanosecond from an unrestrained MD simulation of the CD14 monomer. The exposed surface area drops by ~20% in ~5 ns. C) Sub-panels show slices lengthwise through the center of CD14, with the N-terminal hydrophobic pocket at the top and the C-terminus at the bottom. Waters near the protein are shown as spheres. The left sub-panel is the initial structure (0 ns); the right sub-panel is the structure after 5 ns of simulation time. Waters fill the hydrophobic pocket in the initial structure. By 5 ns, the lid helix (see Figure 6) closes and expels the water from the hydrophobic pocket.

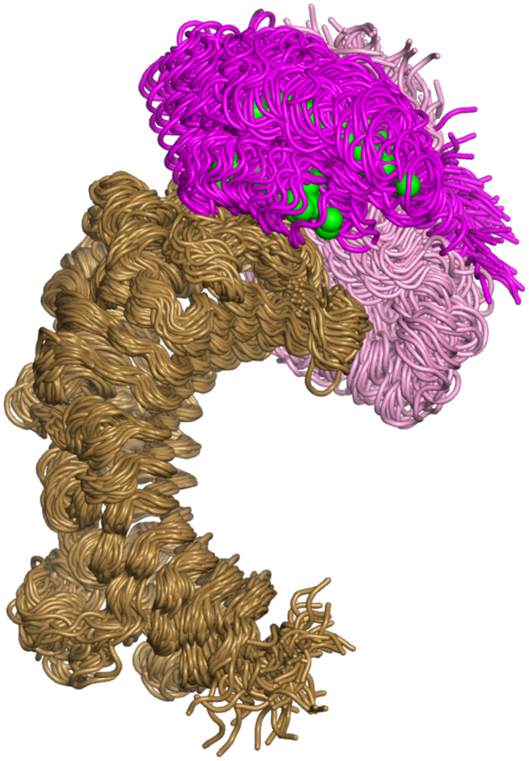

**Figure S3. S100A9 fluctuates relative to CD14 over simulations.** Figure shows overlay of 72 frames extracted from four replicate 500 ns unrestrained MD simulations of CD14 and S100A9, starting from the AlphaFold2 docking model. To reveal the movement of S100A9 relative to CD14, we aligned the core residues of CD14 (residues 100-200) from each frame to the starting structure. The S100A9 dimer is shown in pink (chain A) and magenta (chain B) with calcium ions shown as green spheres. CD14 is shown in brown. We omitted S100A9 residues 1-3 and 91-114 from the figure, as these residues are disordered and obscure the interaction.
